# Identification of 121 variants of honey bee Vitellogenin protein sequences with structural differences at functional sites

**DOI:** 10.1101/2022.01.28.478245

**Authors:** Vilde Leipart, Jane Ludvigsen, Matthew Kent, Simen Sandve, Thu-Hien To, Mariann Árnyasi, Claus D Kreibich, Bjørn Dahle, Gro V. Amdam

**Author notes:** Corresponding Author: Vilde Leipart. Author Contributions Conceptualization: Vilde Leipart, Jane Ludvigsen, Gro V. Amdam Data Curation: Vilde Leipart, Thu-Hien To Formal Analysis: Vilde Leipart Funding Acquisition: Gro V. Amdam Investigation: Vilde Leipart, Jane Ludvigsen, Matthew Kent, Mariann Árnyasi, Thu-Hien To Methodology: Vilde Leipart, Jane Ludvigsen, Matthew Kent, Simen Sandve, Mariann Árnyasi, Thu-Hien To, Claus D Kreibich, Gro V. Amdam Project Administration: Vilde Leipart, Jane Ludvigsen, Gro V. Amdam Resources: Gro V. Adam, Matthew Kent, Simen Sandve, Bjørn Dahle Software: Thu-Hien To Supervision: Jane Ludvigsen, Gro V. Amdam Validation: Vilde Leipart, Jane Ludvigsen, Matthew Kent, Simen Sandve, Mariann Árnyasi, Thu-Hien To, Gro V. Amdam Visualization: Vilde Leipart, Thu-Hien To Writing – Original Draft Preparation: Vilde Leipart, Jane Ludvigsen Writing – Review and Editing: All Authors.

## Abstract

Proteins are under selection to maintain central functions and to accommodate needs that arise in ever-changing environments. The positive selection and neutral drift that preserve functions result in a diversity of protein variants. The amount of diversity differs between proteins: multifunctional or disease-related proteins tend to have fewer variants than proteins involved in some aspects of immunity. Our work focuses on the extensively studied protein Vitellogenin (Vg), which in honey bees *(Apis mellifera)* is multifunctional and highly expressed and plays roles in immunity. Yet, almost nothing is known about the natural variation in the coding sequences of this protein or how amino acid-altering variants might impact structure–function relationships. Here, we map out allelic variation in honey bee Vg using biological samples from 15 countries. The successful barcoded amplicon Nanopore sequencing of 543 bees revealed 121 protein variants, indicating a high level of diversity in Vg. We find that the distribution of non-synonymous single nucleotide polymorphisms (nsSNPs) differs between protein regions with different functions; domains involved in DNA and protein–protein interactions contain fewer nsSNPs than the protein’s lipid binding cavities. We outline how the central functions of the protein can be maintained in different variants and how the variation pattern may inform about selection from pathogens and nutrition.

## Introduction

Protein function relies on the protein’s structural shape, which is dictated by the amino acid sequence that determines the biophysical properties of the molecule. Mutations resulting in non-synonymous single nucleotide polymorphisms (nsSNPs) alter the amino acid sequence and provide an opportunity for new protein variants to enter populations. New variants can be detrimental, neutral, or beneficial in terms of the protein’s impact on phenotype, and these different selective contexts create specific patterns of diversity [1, 2]. For example, multifunctional proteins or proteins at high titers or expressed in several tissues tend to be under strong purifying selection pressure, which results in low diversity [3–5]. An increase in the number of protein-protein interactions is also negatively associated with diversity [6, 7], as is enzyme-function in essential metabolic pathways where changes in the proteins’ active site are unlikely to be beneficial [8]. Conversely, proteins that accommodate diverse or rapidly evolving interaction partners [1, 9]; as exemplified by the histocompatibility complex [10, 11] that recognizes antigens and as observed for membrane- or surface-exposed proteins involved in host-specificity of bacteria [12]. More diversity is also seen in proteins with high designability (i.e., several amino acid configurations accommodate the same fold) [13]. Finally, specific structural features are associated with diversity patterns, such as when exposed structures show more diversity than buried structures [14, 15], or when flexible structures show more diversity than stable β-sheets or α-helices [16].

Vitellogenin (Vg) is a large glycolipo-protein broadly distributed phylogenetically and well known for its role in egg yolk formation. In several species of fish, Vg has immunological functions [17, 18], and in honey bees *(Apis mellifera),* the protein is further recognized for pleiotropic effects on complex behavior [19, 20]. Honey bees are important ecologically and economically as pollinators of native plants and cash crops, and they are key producers of honey, wax, and propolis worldwide [21]. In addition, they represent a flagship species in social insect research [22]. Largely due to these features, Vg has been more intensely studied in honey bees than in most other invertebrates [23]. The protein is found at high titers in hemolymph (insect blood) [24] and localizes to multiple honey bee tissues, including muscle, fat body (functionally analogous to liver and white adipose tissue), gut epithelial cells, and glial cells in the brain [25, 26]. Structurally, the protein has a subdomain of 18 amphipathic α-helices that, together with a β-barrel subdomain and a flexible polyserine linker, form a highly conserved N-terminal domain (ND) [27]. The ND is positioned around a large lipid binding site consisting of a domain of unknown function 1943 (DUF1943) and one β-sheet, followed by a von Willebrand factor (vWF) domain (Figure 1). The final C-terminal region comprises a small structure connected to the vWF domain through a presumed flexible linker [28].

**Figure 1.**
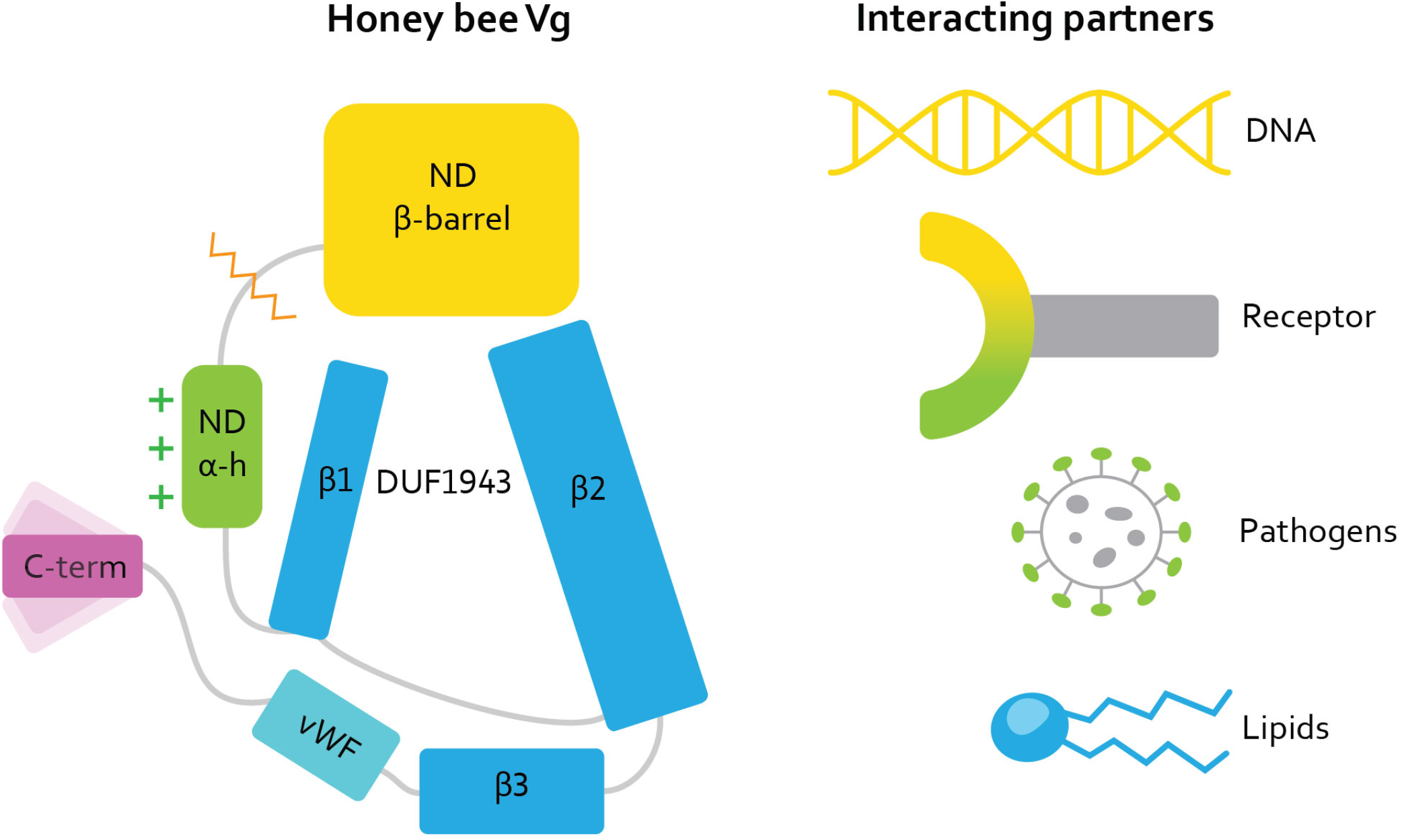
Illustration of the honey bee Vg structure. Vg consists of the N-terminal domain (ND) comprised of two subdomains, β-barrel (yellow) and α-helical (α-h, green), and a lipid binding site (blue), the vWF domain (vWF, cyan), and a C-terminal (C-term, magenta). The orange zig-zag line shows the proteolytic cleavage site on the polyserine linker in ND. The green plus-signs next to the α-helical subdomain illustrate the net positive surface charge. Three β-sheets (β1, β2, and β3) build up the lipid binding site. DUF1943 is defined by β-sheets 1 and 2, while the third sheet is considered part of the lipid binding site; we refer to this structural region as the lipid binding site throughout the article. The C-terminal has been demonstrated to be flexible, as illustrated here. We show the interacting or binding units recognized by honey bee Vg to the right, colored according to the interacting domain or subdomain. We use this coloring scheme throughout the article.

Specifically, the ND likely represents the receptor-binding region of all Vg proteins [29–31]. The ND is also a surface-to-surface contact site in Vg homodimerization, as seen in lamprey *(Ichthyomyzon unicuspis)* [32, 33]. Dimerization at this site is supported in honey bees [28], although Vg appears to be monomeric under most conditions in this insect [34, 35]. Moreover, in honey bees, the β-barrel subdomain of the ND can be proteolytically cleaved at the polyserine linker [36]. The β-barrel appears to subsequently translocate to the nucleus and bind DNA (potentially with co-factors) to influence gene expression [37]. The honey bee ND has a cavity of unknown function in the cleft between the β-barrel and α-helical subdomain [28], while the positively charged α-helical subdomain can account for some of the proteins’ binding to honey bee pathogens [38, 39]. Zooming out, the three structural elements of the large lipid binding cavity create a network of β-sheets with an extensive hydrophobic interior. The hydrophobic core of this site is crucial for the transport and storage of lipids [32], and its structural fold and polarity are conserved across the large lipid transfer protein (LLTP) superfamily to which the Vg proteins belong [40]. The DUF1943 and vWF domains are, in addition, important for innate and mucosal immunity in several species [17, 41, 42]. In contrast, no specific function has been assigned to the C-terminal region of Vg to date [28].

The multifunctionality of honey bee Vg, as well as its high expression, expression in many tissues, and the protein’s interaction with a receptor and dynamics of dimerization may indicate that few Vg variants are found in the bee. The protein’s functions in immunity, in contrast, can suggest that many variants are found. Some support for the latter is provided by previous research [20, 43]. Motivated by these questions, our study seeks a deeper understanding of patterns of variation in honey bee Vg. We examine sequence variation from 15 countries, identify domains under different selective pressures, and characterize the putative functional impact of amino acid changing variants. We reveal 121 unique Vg variants, including 81 nsSNPs that are non-uniformly distributed across the domains and subdomains of the protein. Our analysis illustrates how the structural elements of honey bee Vg experience differing degree of selection pressures.

## Results

### Identification of Vg variants, frequency, and distribution of nsSNPs

Successful amplicon sequencing and variant-calling from 543 individual worker honey bees (diploid females) generated 1,086 full-length *vg* allele sequences, corresponding to 340 unique haplotypes (see Figure S1 for an overview of workflow and Materials and Methods for further details). These haplotypes include different combinations of 81 nsSNPs (see Table S1 for information on the nsSNPs’ properties) resulting in 121 protein variants of honey bee Vg (Table S2; see Figure S2 for an overview of the geographical location of these variants).

In all domains and subdomains of the Vg gene, nsSNPs were identified, with a mean total number of nsSNPs per Vg variant of 5.56 (SD = 1.76). Some nsSNPs occurred more frequently than others: specifically, 15 of the 81 nsSNPs were identified in ≥5 % of the Vg variants. Except for one (p.Arg1292Ser) (6 %), these common nsSNPs caused subtle changes in residue type (Figure 2A). Variants with only common nsSNPs carry the same (one) change in the α-helical subdomain of the ND, and the β-barrel subdomain of the ND and in the C-terminal region typically has few nsSNPs (see Figure 2C for examples). The specific number of nsSNPs and their combinations vary more for the lipid binding site, which thus becomes unique for each Vg variant. In contrast to the common nsSNPs, 20 of 42 (48 %) of the nsSNPs observed only once (i.e., rare nsSNPs) conferred major changes in amino acid characteristics (Figure 2B). For a look at rare nsSNPs, we present a set of Vg variants that includes several rare changes (Figure 2D). In these examples, as seen with the rare Vg nsSNPs overall, we find some in the α-helical subdomain (see variant nr. 34, Figure 2D), and only one change in the β-barrel subdomain, in contrast to several changes in the lipid binding site, including the vWF domain.

**Figure 2.**
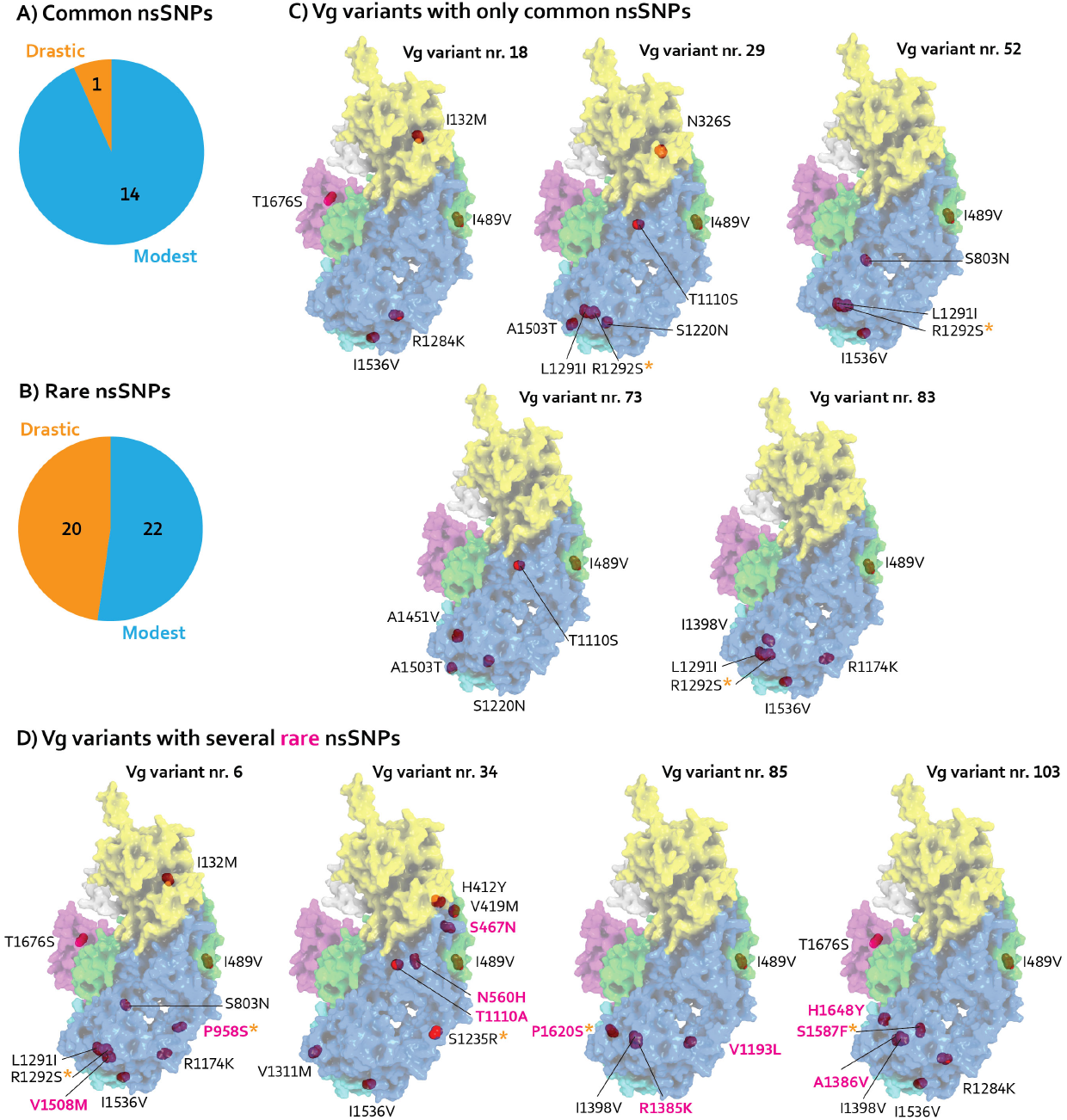
A, B) The common and rare nsSNPs (determined by the number of occurrence in the Vg variants, more than 5 Vg variants are common, while only observed once is rare) are divided by whether they introduce a drastic or modest change in residue type. The drastic substitutions are defined determined by a change of physicochemical properties. For a complete overview of the nsSNPs properties, see Table S1. C) The surface view of five Vg variants (same coloring scheme as in Figure 1) with only common nsSNPs (red spheres). An orange asterisk (*) marks the drastic nsSNPs. D) Vg variants with several rare nsSNPs (labeled in pink) and drastic, labeled as in panel C.

Taken together, the distribution of the rare nsSNPs across protein domains mirrors that of the common nsSNPs. The α-helical subdomain of the ND tends to carry similar changes between variants. In contrast, the lipid binding sites, including the vWF domain, tend to carry more diverse sets of nsSNPs between variants (Figure 3).

**Figure 3.**
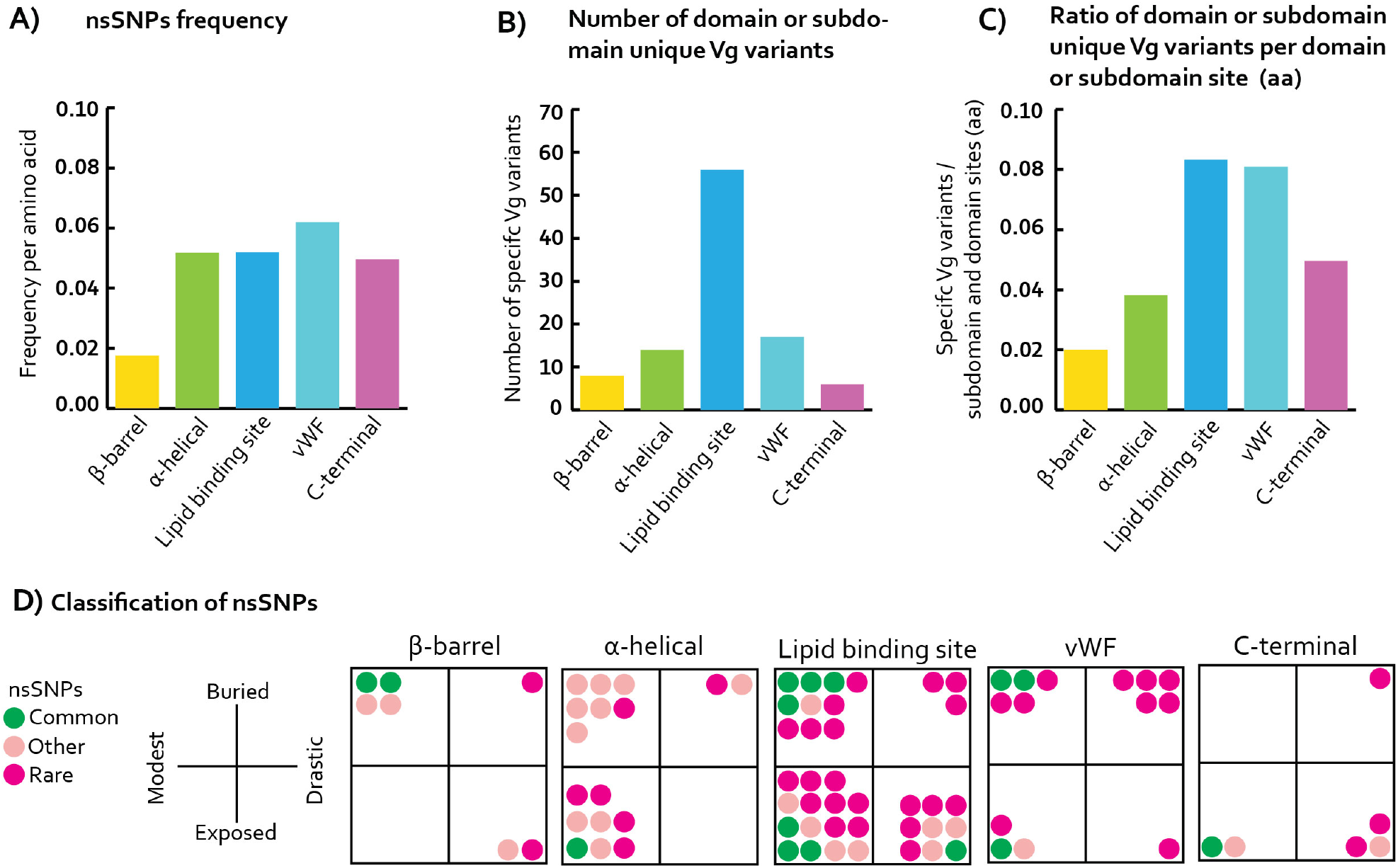
A) The frequency of nsSNPs (used same colors as in Figure 1 for the domains and subdomains) per amino acid (y-axis) presented for the Vg domains and subdomains (x-axis). B) The Vg variants have nsSNPs in different combinations. We divided the Vg variants into domains and subdomains and found the number of unique combinations for the domains and subdomains. These are plotted here (same colors as in panel A). C) The number of unique domains and subdomains used to find the ratio to the size of the domain and subdomain sites (aa). The ratio is plotted here (same colors as in panel A). D) The nsSNPs are colored by how often they were identified on the Vg variants. We considered nsSNPs common when identified on more than 5 Vg variants (green), while the nsSNPs only identified once are considered rare (pink). The nsSNPs identified in 5 to 2 Vg variants were also considered and classified as “other” (light pink). The nsSNPs were divided into the same subdomains or domains used in panels A, B, and C and plotted according to the nsSNPs’ properties (see Table S1 for a complete overview). We calculated the relative solvent accessible surface area (rASA) for each substituted residue, determining how exposed the site is in the protein structure. We considered nsSNPs with a value of 20 % or less as buried; otherwise, they were classified as exposed. The effect of each substitution was determined using a substitution matrix (BLOSUM62) since it shows whether the physicochemical properties are preserved. The nsSNPs with a negative score were considered drastic; otherwise, they were considered modest. We plotted the nsSNPs according to the following classifications: buried or exposed and drastic or modest.

To examine the distribution of 81 nsSNPs in the domains and subdomains, we calculated the frequency of nsSNPs per domain and subdomain site (aa; Figure 3A) and found nsSNP frequency to be lower in the β-barrel subdomain than in the remainder of the domains and subdomains. We subsequently separated the Vg variants into domains and subdomains and counted the number of unique combinations of nsSNPs. This number is higher for the lipid binding site than for the remainder of the domains and subdomains (Figure 3B). The number of amino acids comprising each domain and subdomain varies, which results in a different number of available sites for substitutions at the domains and subdomains. To calculate a ratio to control for this difference, we divided the number of unique Vg variants by the number of sites (aa) per subdomain and domain (Figure 3C). This represents a ratio of unique Vg variant per subdomain and domain. The ratio is higher for the lipid binding site and vWF domain than the remainder of the domains and subdomains (Figure 3C).

We classified the nsSNPs identified in the domains and subdomains into three categories. First, we used the common and rare categories described above and included the remaining nsSNPs (other). Then, we considered if the changes were modest or drastic and calculated whether the substituted residues were at buried or exposed sites in the protein structure. Figure 3D shows the resulting plot for each subdomain and domain. The plot reveals considerable differences between the structural elements of Vg. We assessed whether this variability in distribution and classification of nsSNPs justified a domain- or subdomain-specific approach in the next-step analyses.

### Implications of β-barrel subdomain variants

Only 7 of the 81 nsSNPs were identified in the β-barrel subdomain (Figure 3D), which is less than for other domains (Figure 3A). Except for p.Gly146Ser, all of the nsSNPs cluster at one side of the structure (Figure 4A). Gly146 is buried in the subdomain, close to a set of predicted Zn^2+^-coordinating residues and a proposed DNA binding region (Leipart *et al.* in manuscript)[37]. The remaining nsSNPs increase the polarity of buried residues or increase the hydrophobicity at the surface, except for p.Ile132Met, which maintains the hydrophobic core (Figures 4B and 4C). Overall, the 121 Vg variants identified in this study either contain none or one nsSNP in the β-barrel subdomain, except for Vg variant nr. 5, which carries two common nsSNPs (Figure 4C).

**Figure 4.**
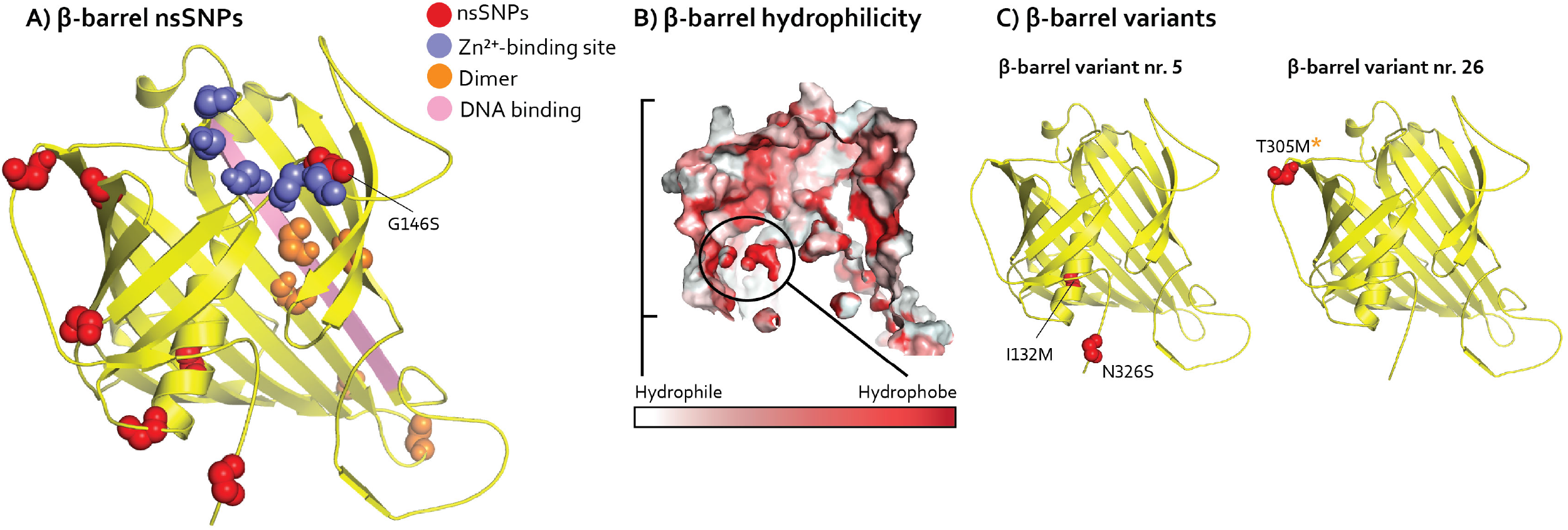
The identified nsSNPs in the β-barrel subdomain (yellow cartoon) are plotted together on the structure, even though the nsSNPs are not identified on the same Vg variant. The spheres represent nsSNPs (red), proposed Zn2+-binding residues (purple) and homodimerization active residues (orange). The DNA binding β-sheet is colored in pink. B) The hydrophobic core adjacent to p.Ile132met is circled, and we show the polar surface for the subdomain. C) β-barrel variant nr. 5 and 26 are shown with the identified nsSNPs (*drastic nsSNPs).

### Implications of α-helical subdomain variants

We identified 17 nsSNPs in the α-helical subdomain (Figure 3D). By mapping the nsSNPs onto the structure, we identified three hotspots of amino acid substitutions (H1, H2, H3; see Figure 5A). The same classification outlined in Figure 3D was repeated here for the nsSNPs in the identified hotspots (Figure 5A). The only common nsSNP (p.Ile489Val, green in the plots in Figures 3D and 5A) is a modest substitution identified in H2. All hotspots contain rare nsSNPs. Out of the 121 Vg variants identified here, 119 variants include one nsSNP in H2, and 15 variants have at least one nsSNP in H1 and/or H3, as shown for Vg variant nr. 24, 41, and 45 (Figure 5B).

**Figure 5.**
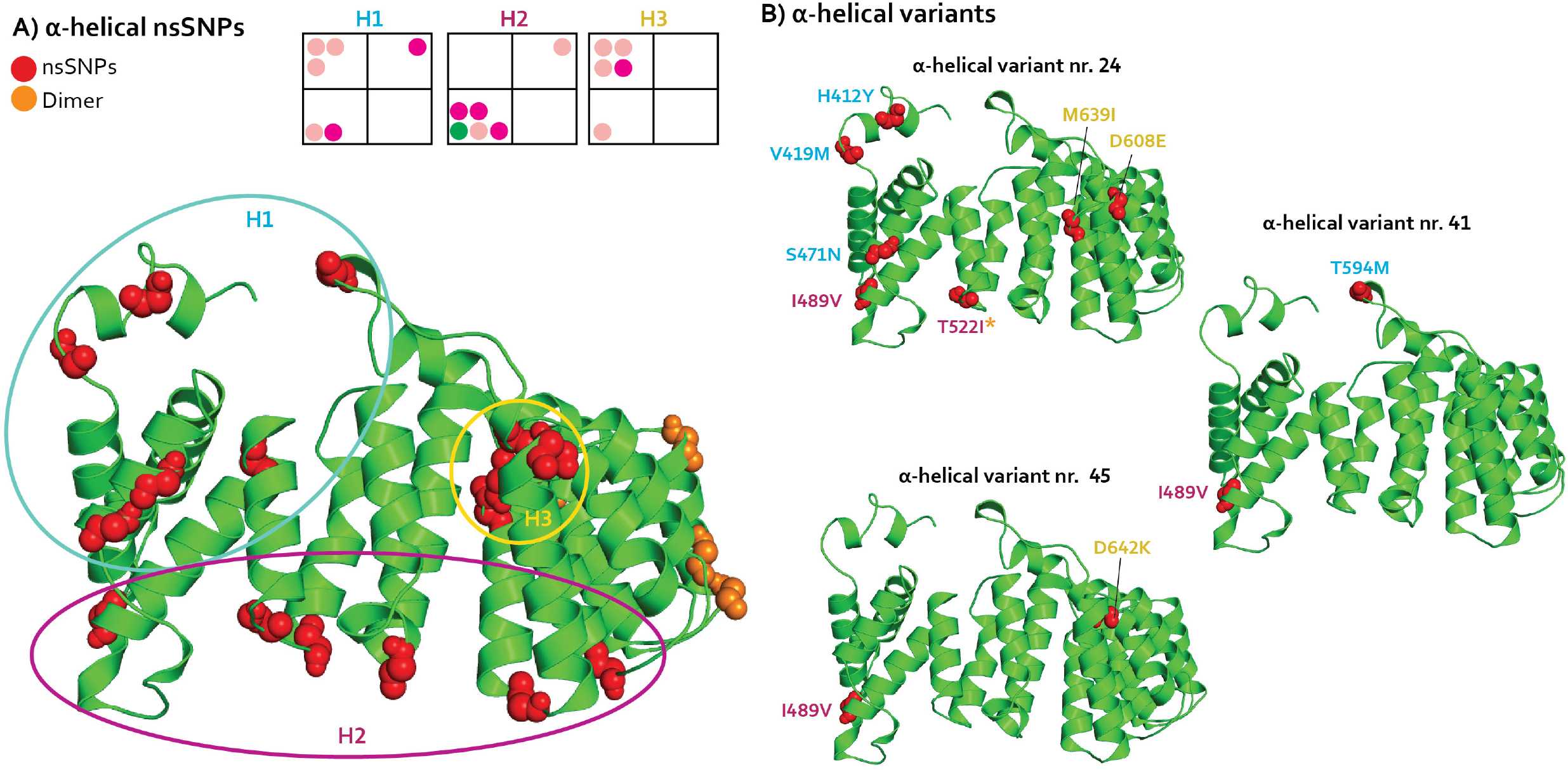
A) The identified nsSNPs in the α-helical subdomain (green cartoon) are plotted together on the structure, even though the nsSNPs are not identified on the same Vg variant. The spheres represent nsSNPs (red) and homodimerization active residues (orange). The identified hotspots H1 (blue), H2 (dark pink), and H3 (yellow) are circled. The nsSNPs are also plotted according to properties per hotspot in the same way as in Figure 3D. B) We show the α-helical variant nr. 24, 41, and 45 with the identified nsSNPs labeled according to the colors of the hotpots used in panel A (*drastic nsSNPs).

Looking at the H1 in more detail, we find that it represents moderate substitutions at three buried and two exposed residues (Figure 5A). The polarity is maintained by these nsSNPs, except for the rare p.Thr594Met, which decreases the polarity of the buried region of the hotspot (see variant nr. 41, Figure 5B). H2 encompasses residues frequently substituted in the short loop regions connecting the α-helices, close to the lipid binding site. These substitutions are modest, except the exposed p.Thr522Ile (see variant nr. 24, Figure 5B). The effects of the nsSNPs on the polarity and electrostatic potential of the structure vary as hydrophobic and hydrophilic residues are introduced. One nsSNP provides a positive charge (p.Asn560His), while another nsSNP removes a negative charge (p.Asp626Asn). The same variability for electrostatic potential is seen in H3, which is buried between two of the subdomain α-helices and the first β-sheet of DUF1943: one nsSNP maintains a negative charge (p.Asp608Glu), while another flips the charge from negative to positive (p.Glu642Lys; see variant nr. 24 and 45, Figure 5B). The remaining three nsSNPs in H3 maintain hydrophobicity at their specific sites.

### Implications of lipid binding site variants

We identified 37 nsSNPs at the lipid binding site (Figure 3D). The nsSNPs were found in 56 combinations (Table S2 and Figure 3B) without discernable clustering into hotspots. The 56 combinations represent a high ratio relative to domain size (aa; Figure 3C). Only three out of the 121 Vg variants lack nsSNPs in the lipid binding site, confirming that it represents a highly diverse protein region. Underlining this level of diversity is the identification of 10 different nsSNPs in just two Vg variants (see variant nr. 1 and 49, Figure 6B).

**Figure 6.**
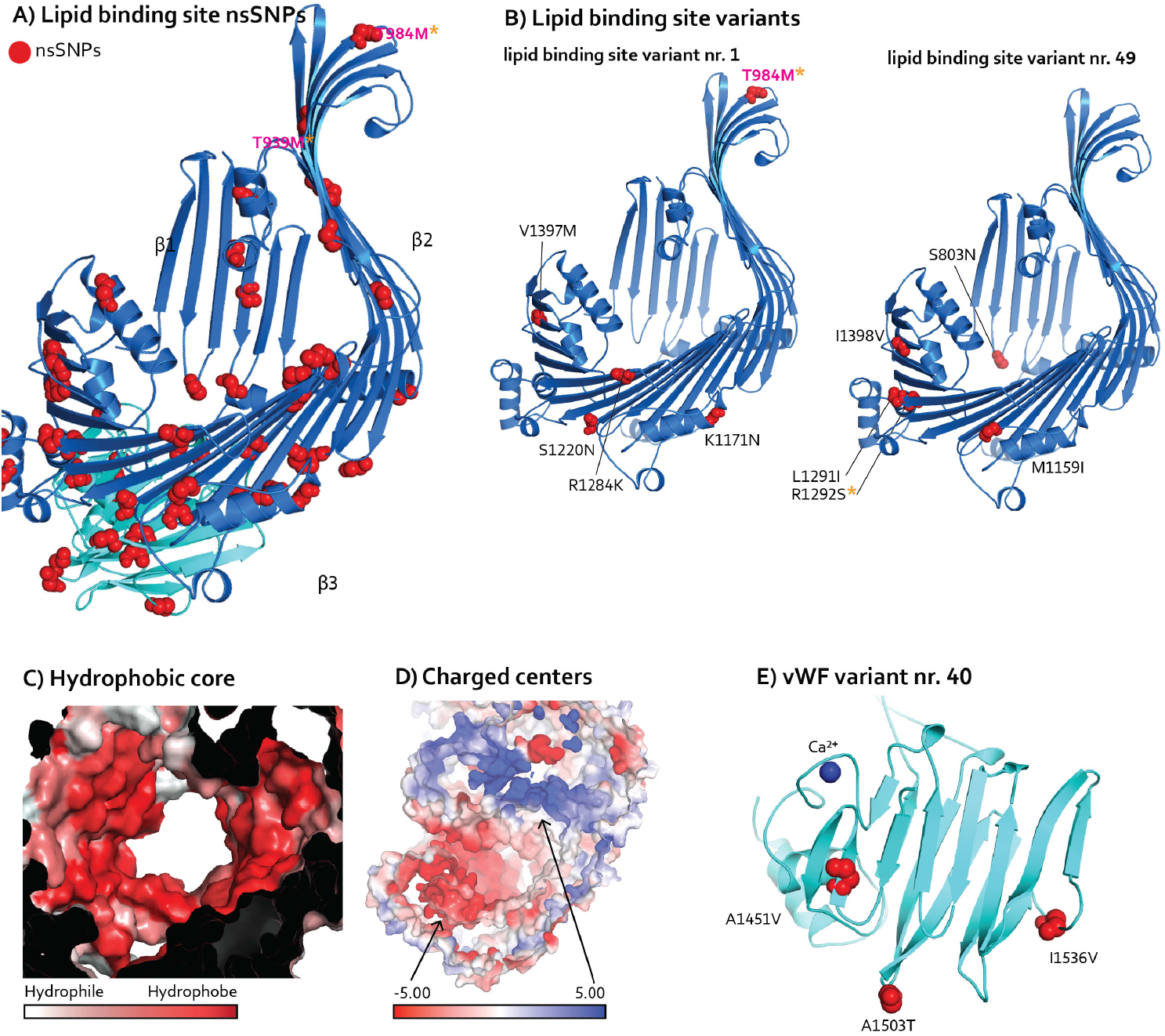
A) The identified nsSNPs in the lipid binding site (blue cartoon) and vWF domain (cyan) are plotted together on the structure, even though the nsSNPs are not identified on the same Vg variant. Spheres represent nsSNPs (red), and the two rare nsSNPs are labeled (pink=rare; *drastic nsSNPs). The three β-sheets shown in Figure 1 are labeled. B) Lipid binding site variants nr. 1 and 49 are shown with the identified nsSNPs (pink=rare; *drastic nsSNPs). C) The lipid cavity is very hydrophobic. D) The two charged centers are shown (black arrows). E) We show vWF variant nr. 40 with the three common nsSNPs. The Ca2+-ion is a blue sphere.

Specifically, drastic substitutions at the lipid binding site were identified at exposed residues, altering the polarity and electrostatic charge of the surface (Figure 3D). This dynamicity of surface residues is a common finding [16], as are moderate substitutions at buried residues [15]. We observed that moderate substitutions do not appear to alter the hydrophobic core or the two charged centers of the Vg lipid binding cavity (Figures 6C and 6D). In addition, however, we find three rare and drastic substitutions at buried residues. Two of these nsSNPs increase the hydrophobicity at the end of the long β-sheet spanning the ND (Figures 6A and 6B), while the third nsSNP increases the polarity of a buried loop, folded away from the domain core.

### Implications of vWF domain variants

We found 14 nsSNPs in the domain (Figure 3D). They were identified in 17 unique combinations, which represents a high ratio relative to domain size (aa; Figure 3C). Overall, the changes are diverse and distributed without discernable hotspots, as we observed for the lipid binding site that interfaces with the vWF domain (Figure 6A).

Interestingly, the vWF domain shows a total of 5 drastic (but rare) substitutions at buried residues. This is the highest number of drastic, buried nsSNPs, compared to the other Vg domains or subdomains (Figure 3D). Three of these nsSNPs either maintain or introduce a polarity, while the other two increase hydrophobicity. Among the 14 nsSNPs in the vWF domain, Ser1587 is the only substituted residue directly exposed to the lipid cavity. This nsSNP introduces a large aromatic residue to the cavity (see variant nr. 103, Figure 2D). Additionally, we find three common nsSNPs that maintain hydrophobicity at buried or exposed sites. These three occur together in Vg variant nr. 40 (Figure 6E). The remaining 5 nsSNPs are modest substitutions. Three are buried and maintain hydrophobicity, while two are exposed and maintain polarity.

### Implications of C-terminal variants

We identified 6 nsSNPs in the C-terminal of Vg (Figure 3D). Four out of the 6 nsSNPs in the exposed structure introduce a serine residue (Figure 7A). These are positioned at the presumed flexible linker or an exposed loop extending from the folded structure, which increases the polarity of the C-terminal.

**Figure 7.**
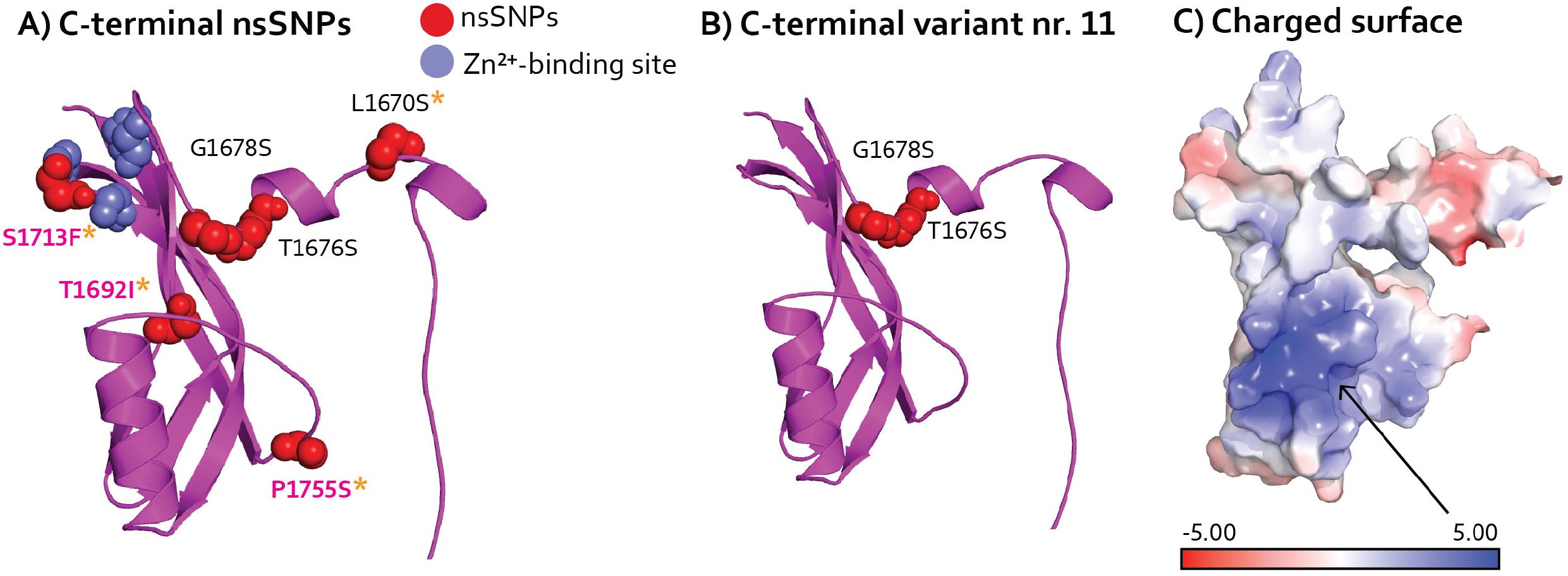
A) The identified nsSNPs in C-terminal (magenta cartoon) are plotted together on the structure, even though they are not identified on the same Vg variant. The spheres represent nsSNPs (red) and proposed Zn2+-binding residues (purple). NsSNPs are labeled (pink=rare; *drastic nsSNPs). B) C-terminal variant nr. 11 is shown with the two serine-introducing nsSNPs. C) The net positive exposed surface is not affected by the nsSNPs.

Two serine-introducing nsSNPs occur together in Vg variant nr. 11 (Figure 7B). The two remaining nsSNPs, not introducing serine, are rare and drastic substitutions (Figure 3D), one increasing the hydrophobicity of the buried structural elements, and the other introducing a large aromatic residue close to a predicted Zn^2+^-binding site (Leipart *et al.* in manuscript). The positive surface charge of the C-terminal is not altered by any of the 6 nsSNPs (Figure 7C).

### Implications of nsSNPs at three domain or subdomain interfaces

Viewing the patterns of nsSNPs in the light of domain or subdomain interfaces, we find that the most variable region of the β-barrel subdomain is adjacent to H1 on the α-helical subdomain. Together, these structures create a hydrophobic and slightly negatively charged cavity (Figures 8A and 8B). A positively charged β-sheet in the DUF1943 domain extends into the cavity, forming an intriguing subdomain interface (Figure 8B). The interface carries 10 nsSNPs: seven introduce a methionine, while the remaining three introduce a tyrosine, leucine, or alanine. One nsSNP decreases the positive charge (p.His412Tyr), while the remaining changes do not influence negative charges of buried or exposed residues, and hydrophobic characteristics are maintained. The conservative nature of these variations is in part explained by the 10 nsSNPs being mostly rare (Figure 8C, for classification of the nsSNPs) and thus unlikely to occur together on one Vg variant (seen in 26 of 121 variants).

**Figure 8.**
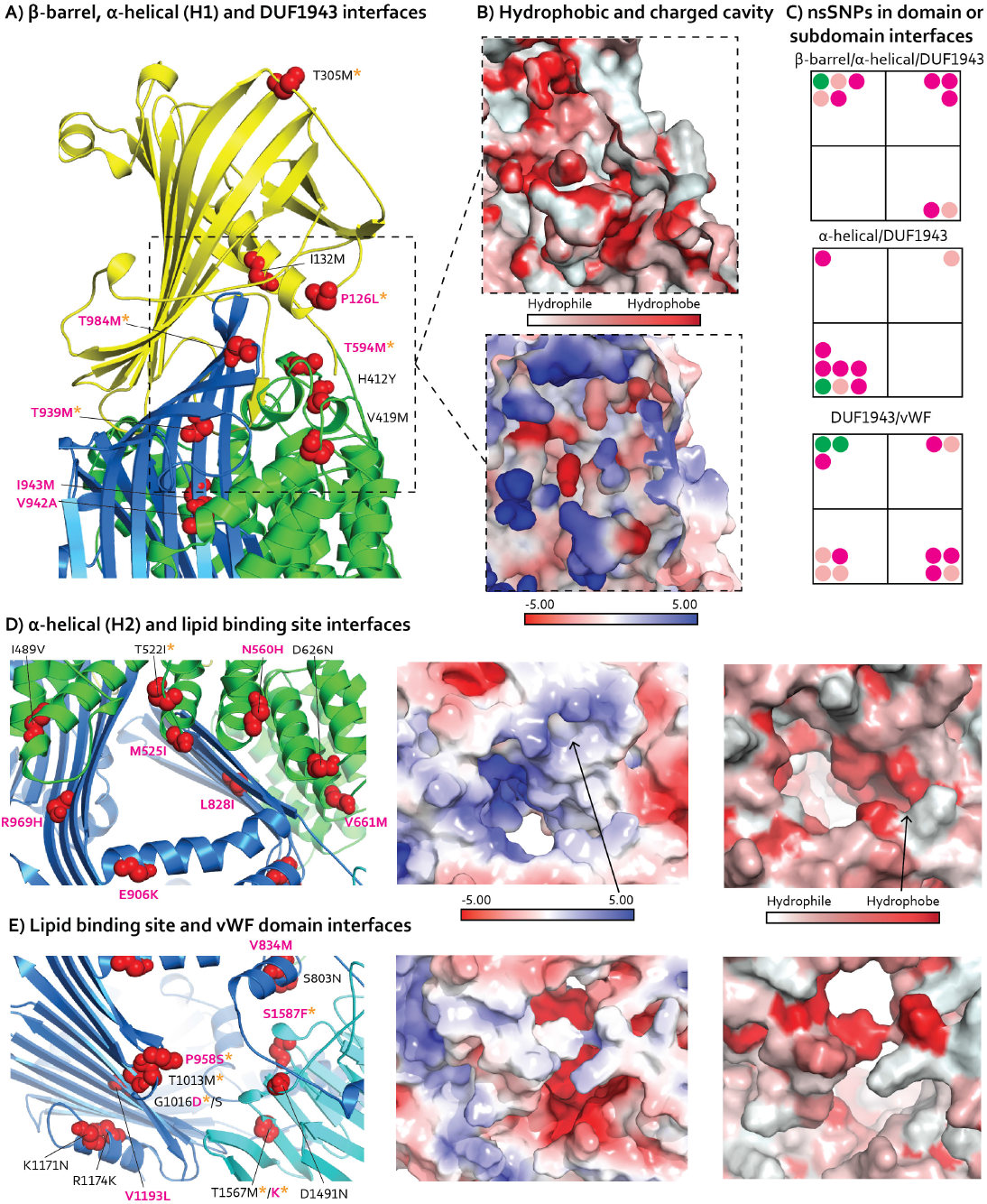
A) The identified nsSNPs in domain or subdomain interface of β-barrel subdomain (yellow), α-helical subdomain (green), and DUF1943 (blue) are plotted together on the structure, even though they are not identified on the same Vg variant. Spheres represent nsSNPs (red) and are labeled (pink=rare; *drastic nsSNPs). B) The hydrophobic core and the electrostatic charges is shown in the dashed boxes, is the same region shown in panel A. C) The same categorization of the nsSNPs identified at three domain or subdomain interfaces, as in Figure 3D. D) The identified nsSNPs in domain interface of H2 in the α-helical subdomain (green) and DUF1943 domain (blue) are plotted together on the structure, even though they are not identified on the same Vg variant. Spheres represent nsSNPs (red) and are labeled (pink=rare; *drastic nsSNPs). We show the positively charged and hydrophobic patches to the right for the same region. E) The identified nsSNPs in the domain interface of DUF1943 (blue) and vWF domain (cyan) are plotted together on the structure, even though they are not identified on the same Vg variant. Spheres represent nsSNPs (red) and are labeled (pink=rare; *drastic nsSNPs). We show the neutral surface to the right for the same region.

Moving on, we find that H2 localizes to an opening where the α-helical subdomain of the ND interfaces with the lipid binding site (Figure 8D). The subdomain interface has a positive charge close to H2, while the edge of the opening (i.e., at the lipid binding site) is hydrophobic (Figure 8D, for classification of the nsSNPs). The nsSNPs in this region are rare and modest substitutions, except for the common p.Ile489Val and the drastic p.Thr522Ile in H2, which maintain and increase hydrophobicity, respectively (Figures 8C and 8D). Two other nsSNPs (p.Asn560His and p.Glu906Lys) slightly increase the positively charged surface, while the hydrophobic region remains undisturbed. The majority of Vg variants identified in this study have only one nsSNP at this subdomain–domain interface (seen in 119 out of 121 variants, including the common p.Ile489Val; excluding this, it is seen in 13 of 121 variants). Next, we observe that the vWF domain is adjacent to an additional opening into the lipid binding site. At this interface, we find 13 nsSNPs (Figure 8E) that do not appear to introduce a consistent type of change. The domain interface is mainly hydrophilic, which is maintained by two common nsSNPs (p.Ser803Asn and p.Arg1174Lys). Other nsSNPs introduce polar and hydrophobic residues: a positive and a negative charge are lost at two different positions (p.Lys1171Asn and p.Asp1491Asn), mirrored by the introduction of a positive and a negative charge at two other positions (p.Gly1016Asp and p.Thr1567Lys). Both buried and exposed residues are modestly or drastically substituted, but these nsSNPs are generally rare (Figure 8C, for classification of the nsSNPs). Adding to the region’s diversity, there are two aa positions with alternative substitutions (p.Gly1016Asp/Ser and p.Thr1567Met/Tyr, see Figure 8E). As observed for the previous domain interface, the Vg variants tend to carry only one nsSNP at the vWF-lipid binding site interface (seen in 36 of 121 variants).

### Implications for the full-length protein structure

When mapping all of the nsSNPs on the surface of the full-length structure of Vg (colored red in Figure 9), we find that the three domain or subdomain interfaces (described above) are located on the same surface side, referred to here as side A (Figure 9). Interestingly, all but one surface-exposed nsSNPs in the ND are located either around the ND cavity where the β-barrel subdomain is interfacing with H1 in the α-helical subdomain or where H2 in the α-helical subdomain interfaces with the DUF1943 domain around an opening to the lipid binding cavity. The one exception is a nsSNP from H3 in the α-helical domain (Figure 9, side A). For the lipid binding site, including the vWF domain, the exposed nsSNPs on side A are found concentrated around a small opening into the lipid cavity, except for two exposed nsSNPs on the vWF domain (Figure 9). Moreover, we find no surface-exposed nsSNPs in the ND when we rotate Vg 180° about the y-axis (as seen in Figure 9, side B). On side B, the exposed nsSNPs are distributed in the lipid binding site, including vWF, making no specific pattern on the surface and not seeming to cluster around the wide opening into the lipid cavity (shaded area in Figure 9, side B). Taken together, our findings demonstrate that honey bee Vg has surface-exposed nsSNPs in every domain and subdomain on side A, while the exposed nsSNPs on side B are only located in the lipid binding site, including vWF.

**Figure 9.**
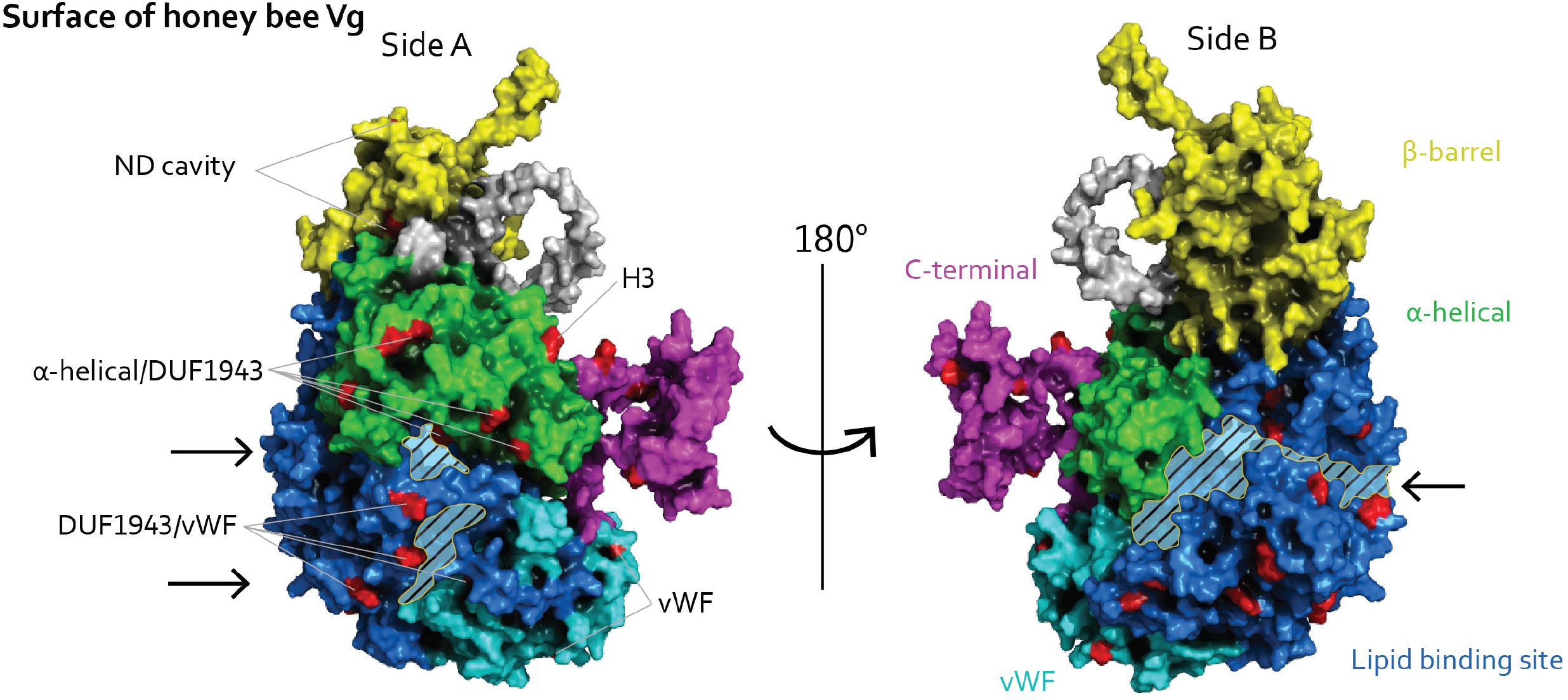
The full-length Vg structure. The colors of domains and subdomains are the same as in Figure 1. Side A: The nsSNPs are colored red on the surface, and the gray lines indicate which domain or subdomain interface the nsSNPs belong to the ND cavity, the α-helical H2 subdomain to the DUF1943, or the DU1943 to the vWF domain. The two smaller cavities (shaded area) leading into the lipid binding site have black arrows pointing to them. Three exposed nsSNPs, not part of a domain or subdomain interface, are marked, one from H3 in the α-helical subdomain and two at the vWF domain. Side B: Rotating 180° about the y-axis reveals a large opening to the lipid binding site (shaded area and black arrow). The surface-exposed nsSNPs are colored red.

## Discussion

This study presents new information on the diversity pattern of Vg. Our geographically broad sampling strategy resulted in over 100 full length Vg protein sequence variants, which is the largest collection of Vg protein variants in any species. Our data confirm the conserved nature of the ND: no changes were observed at aa positions in functional sites for DNA interaction [37] in the β-barrel subdomain, nor at positions suggested for homodimerization at either ND subdomain [28, 33] (see Figures 4A and 5A). The oligomerization state in native honey bee Vg is uncertain [28], but a requirement to protect surface properties involved in homodimerization is supported by our data. Simulating a homodimerization event exposes side A, while ND becomes inaccessible on side B (Figure S3). We find additional support for the conservation of the β-barrel subdomain, since none of the nsSNPs appear to introduce instabilities to the β-barrel fold (Figures 3D and 4A). Similarly, we find evidence supporting studies on varying selection pressures on honey bee Vg. These studies pinpoint the lipid binding site as the primary region of diversity [20, 43], as do our data (see, e.g., Figures 3B and 3D). Yet, in addition to these expected findings, our data reveal information that, combined with the first full-length protein structure for honey bee Vg, contributes to a new understanding of the diversity pattern.

### New insights involving the ND

Given the conserved nature of the ND, our finding of 7 nsSNPs in this region might come as a surprise. As shown, 6 of the 7 nsSNPs cluster at the interface to the α-helical subdomain, adjacent to H1 (Figure 8A). We found that these nsSNPs are rare and tend to introduce hydrophobic residues, particularly methionine. These observations support the idea that selection acts to maintain the characteristics of this structure. Specifically, the region of the 6 clustering nsSNPs is part of a cavity [28], and the conservation of hydrophobic residues is typical for a binding site [44, 45].

The functionality of binding cavities is defined by the residue types, shapes, and locations in the protein [46]. At the β-barrel/α-helical subdomain interface, the β-barrel residues create a hydrophobic and slightly negatively charged region, which meets a positive interior. This structure resembles the large lipid cavity further downstream in the aa sequence. However, the overall shape of ND differs, since the cavity is closer to the protein surface and smaller. We interpret this difference to indicate that the two cavities of honey bee Vg are not functionally equivalent. A distant homolog found in lamprey supports this interpretation, since no phospholipids were observed at the location of the ND cavity [32]. The more conserved nature of the ND cavity compared to the Vg lipid binding site lends further support (Figures 3A–D): the more conserved ND cavity could have a consistent binding partner, while the lipid binding site might interact with various groups of lipids. We suggest that the compatible binding partner of the ND cavity is the Vg receptor. In support of this suggestion, it is assumed that the ND provides the receptor-binding site of the Vg proteins [29–31], and we observe that all but one of the nsSNPs (p.His412Tyr, seen in three Vg variants) introduce no or little change in the electrostatic potential of the ND cavity. Such electrostatic potential is generally important for receptor binding [30, 47]. It has previously been demonstrated that β-sheets in the β-barrel subdomain, as well as α-helices in the α-helical subdomain, have affinity and/or enhanced affinity to the Vg receptor [29–31]. Still, no specific residues in the ND had been specified to participate in this interaction before our work.

The second subdomain in the ND, the α-helical subdomain, has an immune-related function in honey bees that involves the transport of immune elicitors (fragments of bacterial cell wall, i.e., lipopolysaccharides or peptidoglycans) [39] and the recognition of pathogen-associated molecular patterns (PAMP) [38]. The PAMP recognition by Vg is demonstrated for several species of fish [48–50]. High levels of diversity are found in at least some proteins involved in immune defense mechanisms, such as pattern recognition receptors that bind to bacteria via PAMP. In these receptors, the recognition domain is characterized by a leucine-rich repeat that carries nsSNPs, modulating the ability to identify various pathogens [51–53]. Based on this mechanism and the *in vitro* detection of PAMP binding by the α-helical subdomain of honey bee Vg [38], we expected to find a level of diversity in one or more regions of the subdomain. Indeed, we find three nsSNP hotspots: the first (H1) is part of the ND cavity discussed above. The second and third interface with the lipid binding site (H2) or are buried in the subdomain (H3), respectively (see Figures 5A and 8D). In assessing their potential for binding PAMP, we find a high level of diversity at exposed residues in H2. This diversity represents substitutions with a lack of consistency for the introduced residue types that is similarly observed in leucine-rich repeats of protein recognition receptors [51–53]. The buried nature of H3 makes it a less attractive candidate for a direct role in PAMP binding. Instead, nsSNPs could influence subdomain stability and functionality [54–59]. Thus, we speculate that H2 has the potential to be involved in binding specificity with PAMP, while H3 has the potential for being indirectly involved by influencing subdomain functionality for recognition.

Currently, no specific description exists of a molecular mechanism of pathogen binding by the α-helical subdomain of honey bee Vg. Yet, we find 34 positively charged exposed residues (arginine and lysine) that could have an affinity to negatively charged pathogen membrane surfaces [38]. Similar positive surface charge can create high host affinity (but low specificity) for pathogen recognition [60].

Interestingly, the 34 positively charged residues in the α-helical subdomain are conserved in all of the 121 Vg variants identified by our study. We identify no nsSNPs on any exposed 34 arginine or lysine residues (Figure S4). Taken together, the combination of a variable hotspot possibly involved in binding specificity (i.e., H2) and a conserved surface area involved in pathogen affinity (i.e., the 34 positively charged residues) could help provide a molecular understanding of how the α-helical subdomain of Vg contributes to honey bee immunity. At the same time, this insight helps explain why the α-subdomain, overall, may have lower diversity than expected for immune-related activity.

### New insights involving the lipid binding site and vWF domain

The lipid binding site interfaces with H2 and the vWF domain. At both interfaces, we identify nsSNPs that introduce a positively charged residue and a high diversity. Also, when folded, the surface-exposed domain interfaces between the lipid binding site and vWF domain are near the α-helical subdomain (Figure 9A). This structural constellation could imply that the pathogen recognition region of Vg expands beyond the α-helical domain – a proposition supported by previous observation: fulllength honey bee Vg binds PAMP better than the α-subdomain alone [38]. Several members of the LLTP family have similar recognition potential through the α-helical subdomain [40] but have additional protective roles as lipid presenting proteins. For example, the microsomal triglyceride transfer protein (MTP) has an important role in loading endogenous and exogenous lipids onto antigen-presenting cells in the human immune system [61, 62]. Similarly, apolipophorin III and I/II in insects can recognize pathogens [63, 64]. Studies of apolipophorin III show that additional immunological function, such as the ability to regulate and activate hemocytes (immune cells) or stimulate cellular encapsulation, is gained in a lipid-associated state [64]. This conditional functionality is explained by a conformational change when the protein binds lipids [65]. We speculate that the recognition surface presented by honey bee Vg could increase in response to lipid binding; thereby, maintaining the stability of the lipid binding cavity is important for immunological function.

Pathogen membrane surfaces are large relative to a protein [66, 67], so presenting several regions on the protein for affinity and/or specificity is certainly feasible. In this context, we note that the vWF domain of Vg can recognize pathogens in coral *(Euphyllia ancora)* [41] and zebrafish *(Danio rerio)* [17]. Interestingly, we identify a high level of diversity in the honey bee vWF domain (Figure 3C), yet these substitutions mostly occur at buried residues (Figure 3D). The vWF domain is predicted to be an important β-sheet structural region in the lipid binding cavity [28] (Figure 1), and the β-sheet structure is central to the stability of this cavity [40, 57]. Substitutions at buried regions, like those seen for the vWF domain, can affect stability and consequently regulate the size of the lipid load in Vg.

Interestingly, we find exposed residues undergoing changes inside the lipid binding cavity. The lipid cavity interior of Vg is not hypothesized to partake in immune-related activities directly. Instead, the region is recognized for a role in the transport and storage of nutritional phospholipids. Studies of proteins in the LLTP superfamily show that maintaining the large hydrophobic core of the cavity facilitates a high affinity but low specificity for lipid molecules [32, 68]. Our data confirm that the hydrophobicity is conserved in honey bee Vg and suggest that the exposed nsSNPs inside the lipid cavity might influence lipid specificity. Phospholipids usually occupy the positively charged center, as shown in the lipid cavity for a distant homolog [32]. We find diversity at regions close to this charged center, suggesting that phospholipids might enter the cavity here (side A, Figure 9A). These diverse regions might also influence specificity for lipid molecules as well as pathogen specificity, as discussed above. Thus, overall, an evolutionary arms race with changing pathogens that further vary at different geographies could be a possible explanation for the pattern we observe, as suggested in previous research [69–71].

### New insights involving the C-terminal region

We confirm the C-terminal region on honey bee Vg to be soluble and find 4 nsSNPs introducing polar residues (Figure 7A, seen in 35 Vg variants). This finding supports our previous study showing the region is exposed and connected to a presumed flexible linker [28]. We additionally provide new evidence showing a conserved positively charged surface (Figure 7C). A positively charged C-terminal region in other proteins has been linked to signaling for recruitment and translocation [72], protein assembly [73], and sensing changes in the extracellular environment [74]. Honey bee Vg has been demonstrated to sense oxidative stress [75] and suggested protecting honey bees from reactive oxidative species. Our earlier study shows that two disulfide bridges are conserved in the C-terminal region, which is proposed to coordinate Zn^2+^ (Leipart et al. 2021 *in manuscript)* (Figure 7A). Proteins with a positive surface charge and disulfide bridges on neighboring residues, sometimes including Zn^2+^, are shown to protect against oxidative stress [76, 77]. Our findings support a conserved polarity and positive charged region; thus, we speculate that the C-terminal has a similar functional role.

### Concluding remarks

None of the nsSNPs identified here are detrimental for honey bee Vg. The structural fold in the ND is highly conserved, and the drastic changes in the remaining domains are either exposed at the surface or buried at non-structural loop regions, except for the p.Thr939Met shown in Figure 6A. These nsSNPs increase the hydrophobicity at the protein core, which is unlikely to reduce structural stability. All of these observations are expected for a protein that is essential for fitness in its yolk-precursor role. At the same time, we observe new variability patterns that are likely associated with aspects of lipid binding. In assessing these nsSNPs, we provide new insights on the possible interface between Vg, its lipid cargo, and honey bee pathogens. We believe these suggestive findings are thought-provoking and warrant further study. Additionally, it is worth mentioning that the long-read sequencing technology used here creates an opportunity to identify and characterize genomic structural variants that are difficult or impossible to detect with alternative approaches [78, 79]. Such variants can significantly impact protein structure and should be receiving increasing attention in studies seeking to link genotype to phenotypic variation. Correspondingly, a preliminary examination of our data suggests the presence of larger structural variants (deletions) that will be fully explored in a future manuscript.

## Materials and Methods

### Bee sampling

452 samples of *Apis mellifera* were collected from Europe. Nine protected *Apis mellifera mellifera* apiaries were selected and sampled based on earlier introgression studies [69, 71]: Norway (Flekkefjord, N=30; Rena, N=32), Sweden (Jämtland, N=30), Denmark (Læsø, N=32), Scotland (Isle of Colonsay, N=30), Ireland (Connemara, N=30), Poland (Augustów Primeval Forest, N=30), the Netherlands (Texel, N=30), and France (Les Belleville, N=30). Samples from six European subspecies, from separate apiaries, were chosen for comparison: Slovenia (*A. m. carnica,* N=25), Italy (*A. m. ligustica,* N=30), Portugal (*A. m. iberiensis,* N=30), Macedonia (*A. m. macedonica,* N=33), Malta (*A. m. ruttneri,* N=30), and Turkey (*A. m. anatolica,* N=30). The samples from Europe were provided by researchers and managers of breeding associations working with each subspecies to ensure that samples were obtained from purebred populations. In addition, we collected 186 samples from the USA, used as one control group, from 6 different apiaries covering the north, west, south, northeast, east, and central regions: Minnesota (N=33), California (N=30), Arizona (N=30), Maryland (N=30), North-Carolina (N=33), and Illinois (N=30), respectively. To ensure genetic variation among the samples, the collectors in Europe and the USA sampled 25 to 33 bees from three to six separate hives in their apiaries. The specimens were collected and shipped in 2 ml Eppendorf tubes filled with 1.9 ml 96 % ethanol and stored at −20 °C.

### gDNA extraction

Genomic DNA (gDNA) was extracted from the thorax of each bee. The head, wings, legs, and abdomen were removed, before the thorax was washed in PBS for 5 minutes. The equipment used for dissection was washed in 10 % chlorine and 96 % ethanol between every bee. After washing, the thorax was cut in half vertically and weighed, with weights ranging from 18 to 30 mg. Half of each thorax was used in the DNA extraction protocol. The thorax piece was placed in a tube filled with 200 μl ATL buffer (1:2 ratio) and three sterile ceramic beads (2.8 mm). The samples were ground in Retsch® mixer mill MM 400 (Retsch GmbH, Germany) at 15/s for 20 seconds, before 20 μl Proteinase K and 2 μl Rnase A were added and mixed by vortexing, and the samples were incubated at 56 °C overnight while mixing. The remaining steps followed the QIAGEN® DNeasy® Blood & Tissue Kit standard protocol (QIAGEN, Redwood City, CA). The eluate was eluted twice with a final volume of 100 μl. The concentration was measured on Qubit® 2.0 Fluorometer using the Qubit™ dsDNA HS Assay kit standard protocol (ThermoFisher Scientific, Waltham, MA). The extracted gDNA was run on 0.4 % TAE Agarose gels containing TAE buffer containing StainIN™ GREEN Nucleic Acid Stain (highQu, Germany), at 40V for 1h and 50 min, with the Thermo Scientific™ GeneRuler™ High Range DNA ladder to determine the size and quality of gDNA. Eluted gDNA was stored at −20 °C for 1–2 days, then at −80 °C.

### PCR, pooling, and clean-up

To enable the simultaneous sequencing of amplicons from 543 bee samples, a two-tier barcoding strategy was used, whereby barcodes were included in both the PCR primers and the sequencing adapters. PCR primers were developed to amplify the full-length *vg* gene (including introns) from position 5,029,433 to 5,035,683 in NC_037641.1 [80] (see Table S3 for the primer sequences). In addition to the *vg*-specific sequence, unique barcodes from the PCR Barcoding Expansion 1-96 kit (EXP-PBC096; Oxford Nanopore Technologies, see Table S3 for barcode sequences) were incorporated into the 5’ ends of the forward (n=8) and reverse (n=12) primers, which enabled 96 different barcode combinations. PCR was performed in 96-well plates, wherein each PCR reaction contained 10 ng gDNA, a unique combination of forward and reverse primers (0.5 uM each), 0.5U Q5® High-Fidelity DNA Polymerase (New England BioLabs, MA, USA), 1X Q5 Reaction Buffer, 200 μM dNTP, and Nuclease-free water, to a final volume of 25 μl. Cycling conditions were as follows: 98 °C for 1 min, 30 cycles of 98 °C for 10 s, 58 °C for 30 s, 72 °C for 5 min, and then 72 °C for 7 min and a hold at 4 °C. One positive control sample and one negative control (PCR water) were included for each of the 6 PCR plates that were run. After PCR, the concentration of each amplicon was measured in a plate reader using PicoGreen (ThermoFisher Scientific, Waltham, MA). The positive and negative controls were checked on a 1 % TAE agarose gel to verify amplification and the lack of contamination (see Figure S5 for agarose gel). From each of the 94 samples within each plate, 16 ng was pooled, creating six plate pools (see Table S3 for a plate set up used for each pool). The six plate pools had concentrations ranging from 5.4 to 10.8 ng/μl (Qubit® 2.0, dsDNA BR Assay) and volumes ranging from 392.4 to 731.4 μl. Each pool was concentrated and purified using 0.75X AMPure XP beads (Beckman Coulter, Brea, CA) before being eluted in 60 μl nuclease-free water pre-heated to 50 °C. The concentration of each pool was measured (Qubit® 2.0, dsDNA BR Assay) and found to range from 10.9 to 22.8 ng/μl. Three of the pools with concentrations lower than 15 ng/μl were up-concentrated using a vacuum centrifuge to be able to start with a minimum input of 620 ng amplicons from each pool.

### Library preparation and Nanopore sequencing

For Nanopore sequencing, the library was prepared using the Ligation Sequencing kit 1D (SQK-LSK109) and the Native Barcode Expansion kit (EXP-NBD104), following the “Native barcoding amplicons” Nanopore protocol. The workflow is illustrated in Figure S1A. Briefly, 620 to 850 ng amplicons from each plate pool were used as input to prepare the DNA ends for barcode attachments; native barcodes NB01-NB06 were then ligated to the end-prepared amplicons. After measuring the concentration of the six native barcoded sample plate pools, equal amounts from each pool were combined, and a total of 800 ng mix was taken to adapter ligation. After flow cell priming, 200 ng (equal to 50fmol) final prepared library was loaded into a PromethION flow cell (v9.4.1). MinKNOW v20.06.18 was used for operating sequencing. Base-calling and filtering were performed with Guppy v4.0.11 using the “High-accuracy sequencing” base called model, and the minimum qscore for read filtering was 7. Oxford Nanopore Technologies sequence data were base called real-time using the MinKNOW Fast base calling model from Fast5 into FastQ file format. Raw reads were classed as passed by MinKNOW based on the average read quality score >7.

### Bioinformatic pipeline

The bioinformatic pipeline is illustrated in Figure S1B. About 18 million raw reads were downloaded from the PromethION sever and demultiplexed each native and inner barcodes into separate samples using cutadapt v. >=2.10 [81]. The error rate for the inner barcodes was set to 0.17, and the minimum and maximum length of reads after trimming the inner barcodes was set to 6,000 and 7,000, respectively, reducing the number of raw reads to 6,193,310. Each read was written into a separate folder, and the native and inner barcodes and primer sequences were removed from the reads. The medaka tool (v. 1.0.3 https://nanoporetech.github.io/medaka/index.html, source code, and analysis scripts (available at https://github.com/nanoporetech/medaka) were used to create consensus sequences and variant calling. A consensus sequence for each demultiplexed sample was generated using medaka_consensus based on reference sequence NC_037641.1 [80]. To create haplotype consensus sequences, the phased alignments of the medaka_variant pipeline were first applied and separated the reads into haplotypes for each sample. The medaka_consensus was then re-used, with the same reference sequence as above, to generate a consensus sequence for each haplotype. The variant calling pipeline of medaka was also used for SNP calling for each haplotype using the same reference sequence. The pipeline was implemented using snakemake v.>=5.6.0 (available at https://gitlab.com/cigene/computational/bee_amplicon). We illustrate the pipeline in Figure S1B. The downstream analysis was done on the allele sequences generated from a minimum of 100 raw reads (31 samples had fewer than 100 reads and were not included in the downstream protocol). This resulted in 1,086 allele sequences, generated from an average of 6,497.34 (SD=5,328.55) raw reads per allele sequence.

### Identifying Vitellogenin variants

The raw allele sequences were uploaded to Geneious Prime v.2019.0.03, where we created FASTA files starting at first to the last codon for the *vg* gene (6109 bp, including introns, NP_001011578.1). DNA Sequence Polymorphism v.6.12.03 [82] was used to identify 340 haplotypes and the 81 nsSNPs (See table S1 for an overview of the nsSNPs properties). The nsSNPs are written using the Human Genome Variation Society [83]. Haplotypes with identical nsSNPs combinations were identified as identical Vg variants. The Vg variants are presented in Table S2. The AlphaFold prediction of full-length honey bee Vg was generated from UniProt ID Q868N5, and we used this sequence as a reference for nsSNP analysis.

### Structural analysis

The structural analysis was performed in PyMol v.2.4.1 [84] using AlphaFold Vg structure [28]. We considered nsSNPs identified in more than 5 Vg variants as common and identified only one as rare. Other nsSNPs identified in 5 to 2 Vg variants were also considered and classified as “other.” The relative solvent accessible surface area (rASA) was calculated in PyMol, and residues scoring <20 % were deemed buried [85]; otherwise, they were classified as exposed, although thresholds from 5–25% have been used in literature. The rASA calculation indicates how exposed the residue is at the specific position in the protein structure [86]. The similarity between amino acids was classified for each substitution using a substitution matrix [87]. A negative score indicates that the physiochemical properties are not preserved. Negative scores in the BLOSUM62 matrix were considered drastic; otherwise, they were considered modest. We illustrate these three characteristics for each nsSNP in Figures 3D, 5A, and 8C. The Eisenberg hydrophobicity scale [88] was used to analyze hydrophobicity. The APBS electrostatic plugin in PyMol was used to identify charged regions, and the illustrations were made in PyMol.

## Abbreviations

(DUF1943): The domain of unknown function 1943,
(H): hotspot,
(LLTP): large lipid transfer protein,
(ND): N-terminal domain,
(nsSNPs): non-synonymous single nucleotide polymorphisms,
(PAMP): pathogen-associated molecular patterns,
(rASA): relative solvent accessible surface area,
(Vg): Vitellogenin,
(vWF) domain: von Willebrand factor

## Acknowledgments

We extend our greatest gratitude to the researchers and managers of breeding associations who sampled, handled, and shipped the bee samples collected for our research herein: Anja Laupstad Vatland (Managing Director at Molti AS, Norway), Tor Erik Rødsdalen (Leader of Norsk brunbielag), Ingvard Arvidsson (Adviser at Nordbiföreningen, Sweden), Flemming Vejsnæs (Adviser at Danish Beekeepers Association), Andrew Abrahams (Manager of Colonsay Black Bee Reserve, Scotland), Gerard Coyne (Vice Chairperson at The Native Irish Honey Bee Society and Regional Director of Connacht), Małgorzata Bieńkowska (Lab head at the Research Institute of Horticulture in Skierniewice, Poland), Romée van der Zee (Dutch Center for Bee Research), Klébert Silvestre (President of the Center for Technical Apicultural Studies of Savoie, France), Peter Kozmus (Professional Leader of Breeding Program for Carniolan Honeybee for the Slovenian Beekeepers’ Associations), Cecilia Costa (Researcher at Council for Agriculture Research and Agricultural Economy Analysis, Bologna, Italy), Maria Alice de Silva Pinto (Coordinator Professor at Instituto Politécnico de Bragança, Portugal), Aleksandar Uzunov (Associate Professor at Faculty of Agricultural Sciences and Food, Skopje, North Macedonia), Thomas Galea (committee member of the Malta Beekeepers Association), Irfan Kandemir (Professor at Department of Biology, Ankara University, Turkey), Adam G. Dolezal (Assistant Professor – Entomology at School of Integrative Biology, University of Illinois), Olav Rueppell (Florence Schaeffer Distinguished Professor of Science at Department of Biology, University of North Carolina at Greensboro), Jay Evans (Research Entomologist at Bee Research Laboratory, United States Department of Agriculture, Maryland), Tim Kenney (Beekeeper and manager of Red Mountain Cattle Company, Arizona), Randy Oliver (Manager of Scientific Beekeeping, California), Marla Spivak (Professor in Entomology, University of Minnesota), and Mike Goblirsch (Post-doc at the Spivak Honey Bee Lab, University of Minnesota). We thank you all for your cooperation. The authors acknowledge The Research Council of Norway grant number 262137 for funding toward running costs and positions and BioCat (RCN grant number 249023) for travel grants and conference support.

The authors declare no conflicts of interest.

## Supporting Information

**Figure S1.**
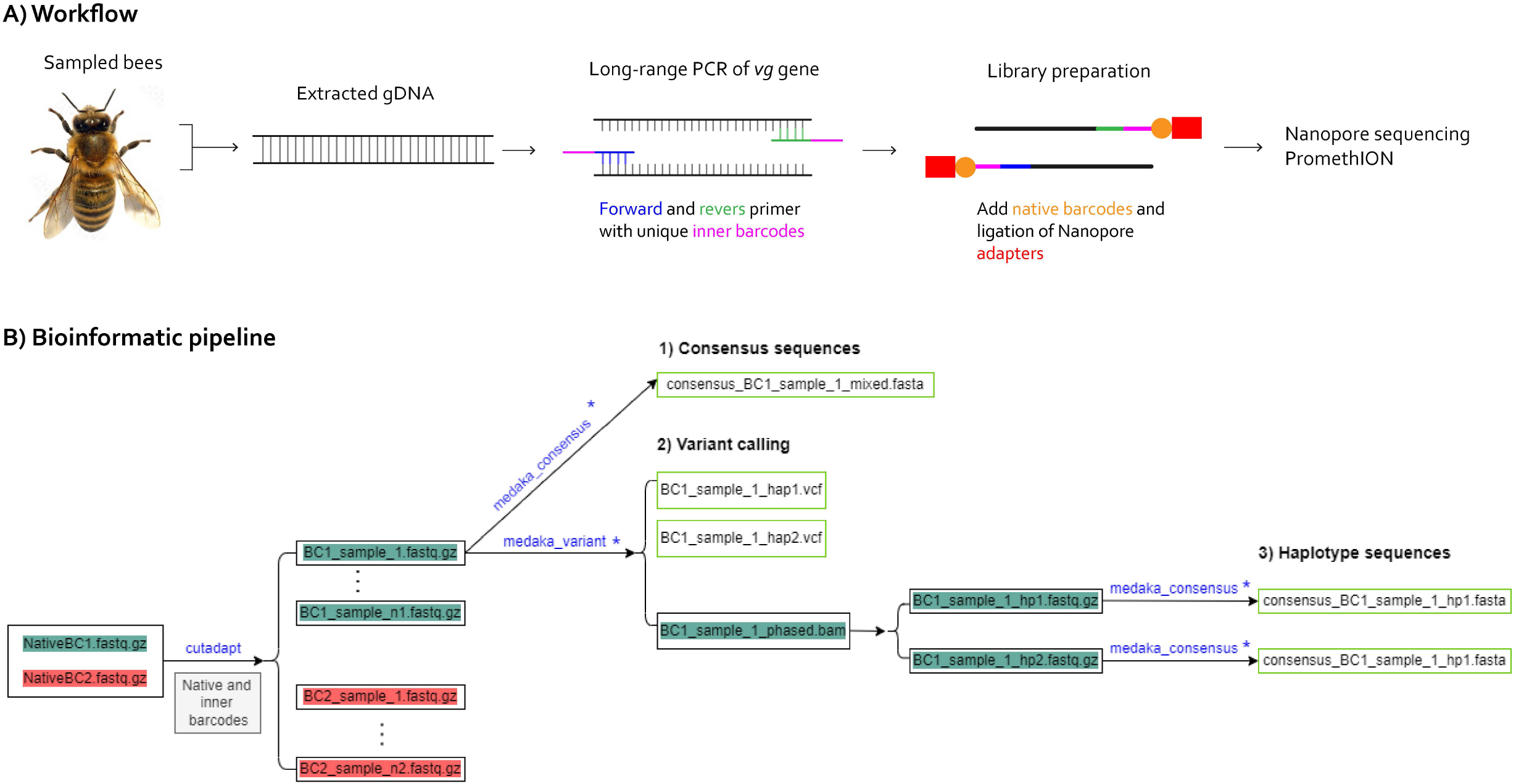
A) Illustration of the workflow. The thorax of the sampled bees was used to extract gDNA. The *vg* gene (6109 bp) was amplified using barcoded primers (see Table S3 for the primer and barcode sequences). We used long-range PCR to amplify the full gene. A native barcode and adapter were ligated to each amplicon sequence before being loaded onto PromethION for Nanopore sequencing. B) Overview of the bioinformatic pipeline. The tools used at each step are written in blue letters above the arrow. First, the raw sequences (first box, NativeBC) were sorted using the native and inner barcodes (see Table S3), resulting in a long list of raw sequences for each sample (second box, sample). For each sample set, three tools were used to generate 1) a consensus sequence for each sample, 2) variant calling files, and 3) haplotype sequences. The blue asterisk (*) at these three steps illustrates that the reference sequence (NC_037641.1 [80]) was applied. (TIFF)

**Figure S2.**
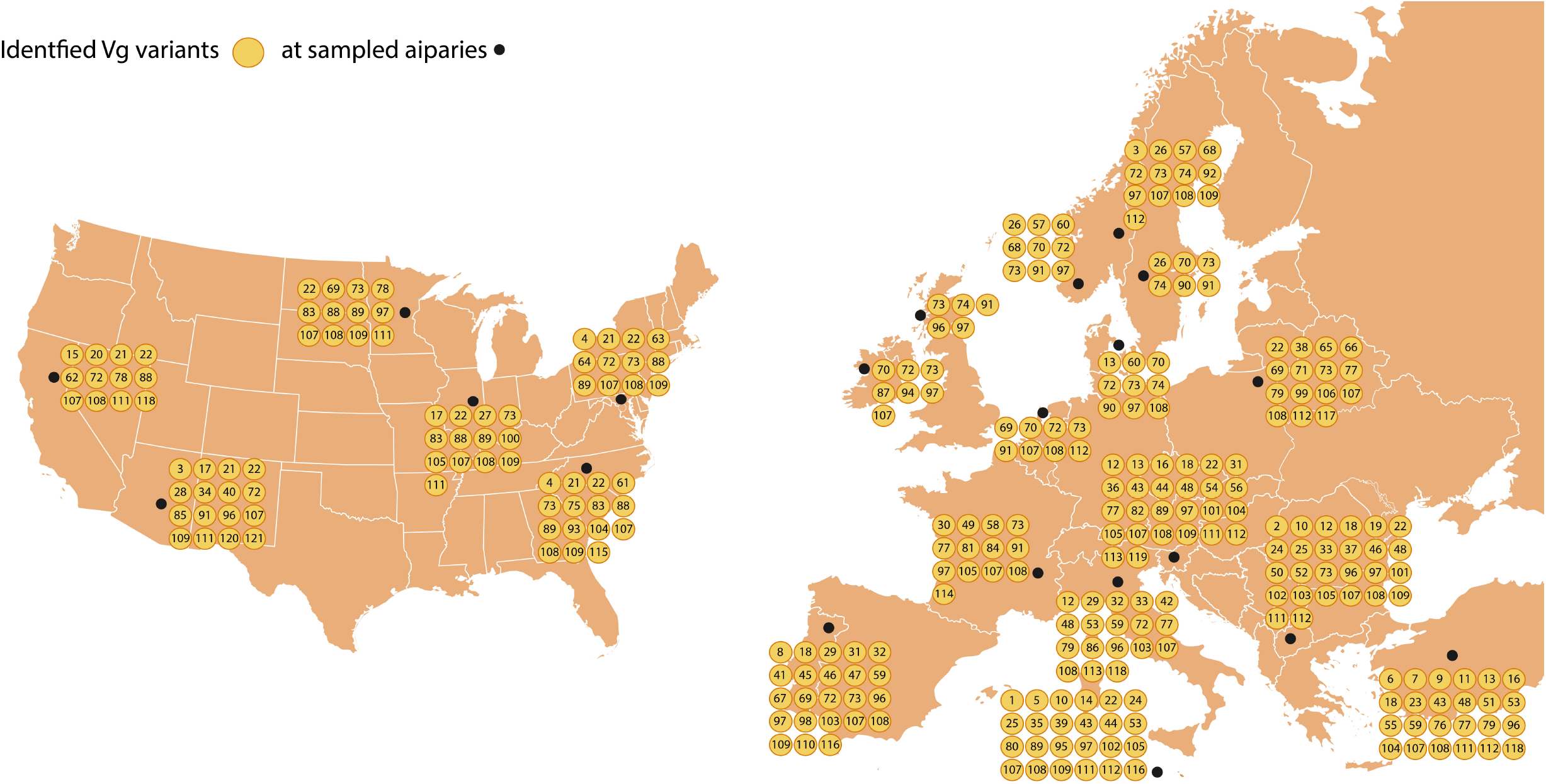
The identified Vg variants (yellow circles, numerated from Table S2) are plotted according to their geographical location. The sampled populations are shown as black dots (more information about the sampled locations is provided in the first section in “Materials and Methods”). (TIFF)

**Figure S3.**
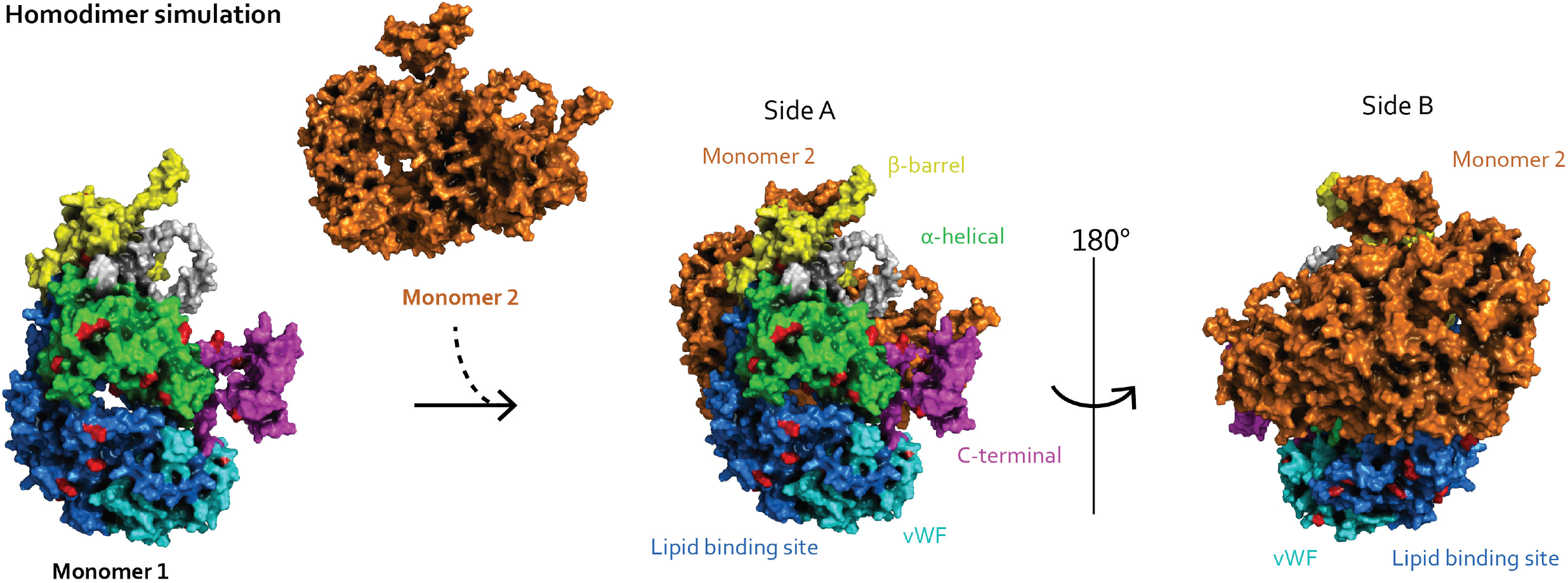
Simulation of the proposed homodimerization event. The full-length Vg is colored the same way as in Figure 9 and represents monomer 1. The second monomer is colored orange. In the event of dimerization, side A will still be exposed, while the ND could be shielded by monomer 2 on side B. (TIFF)

**Figure S4.**
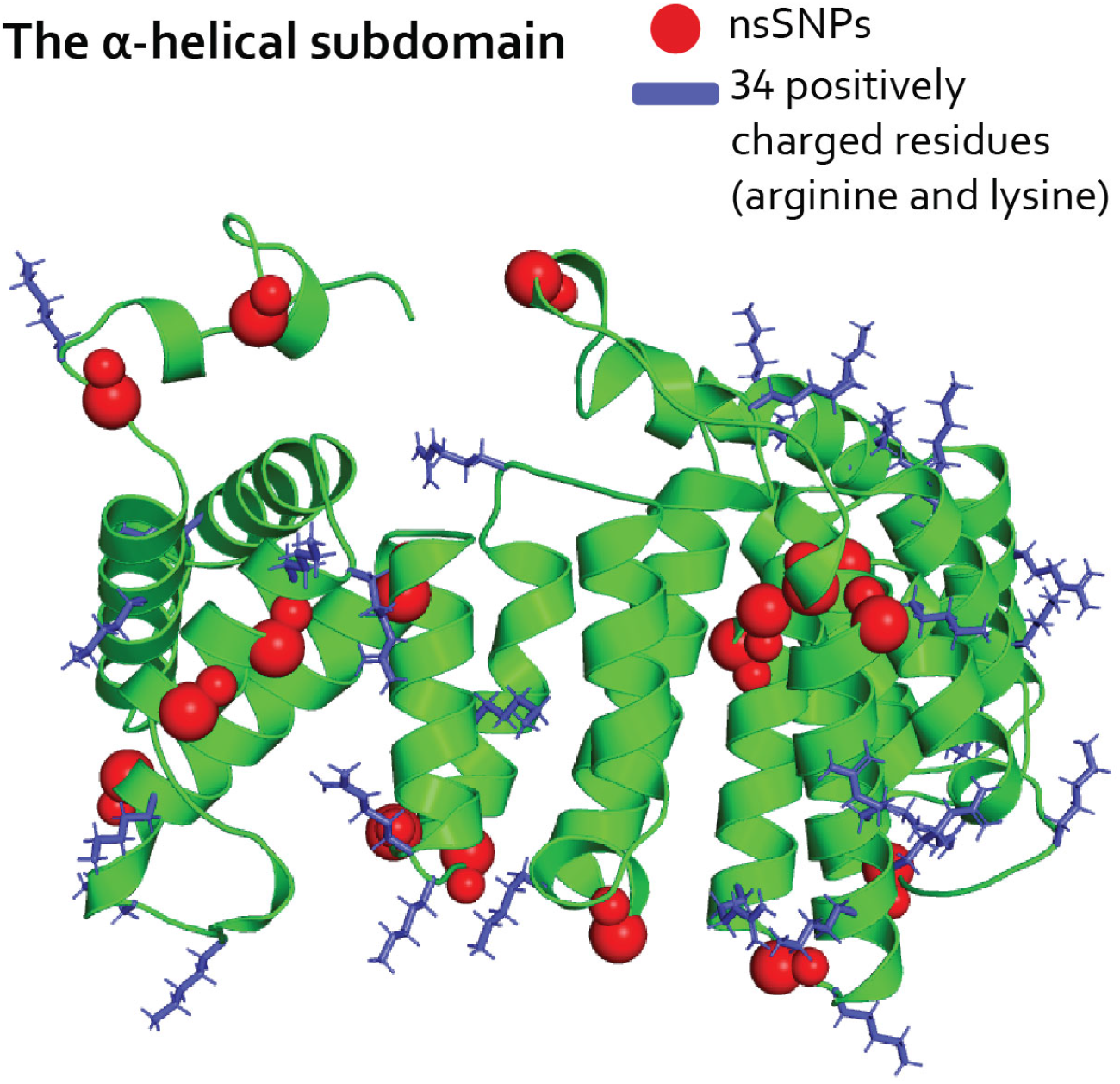
The α-helical subdomain is green with the nsSNPs (red spheres) and the 34 positively charged residues as blue sticks. No changes are introduced at the positively charged residues. (TIFF)

**Figure S5.**
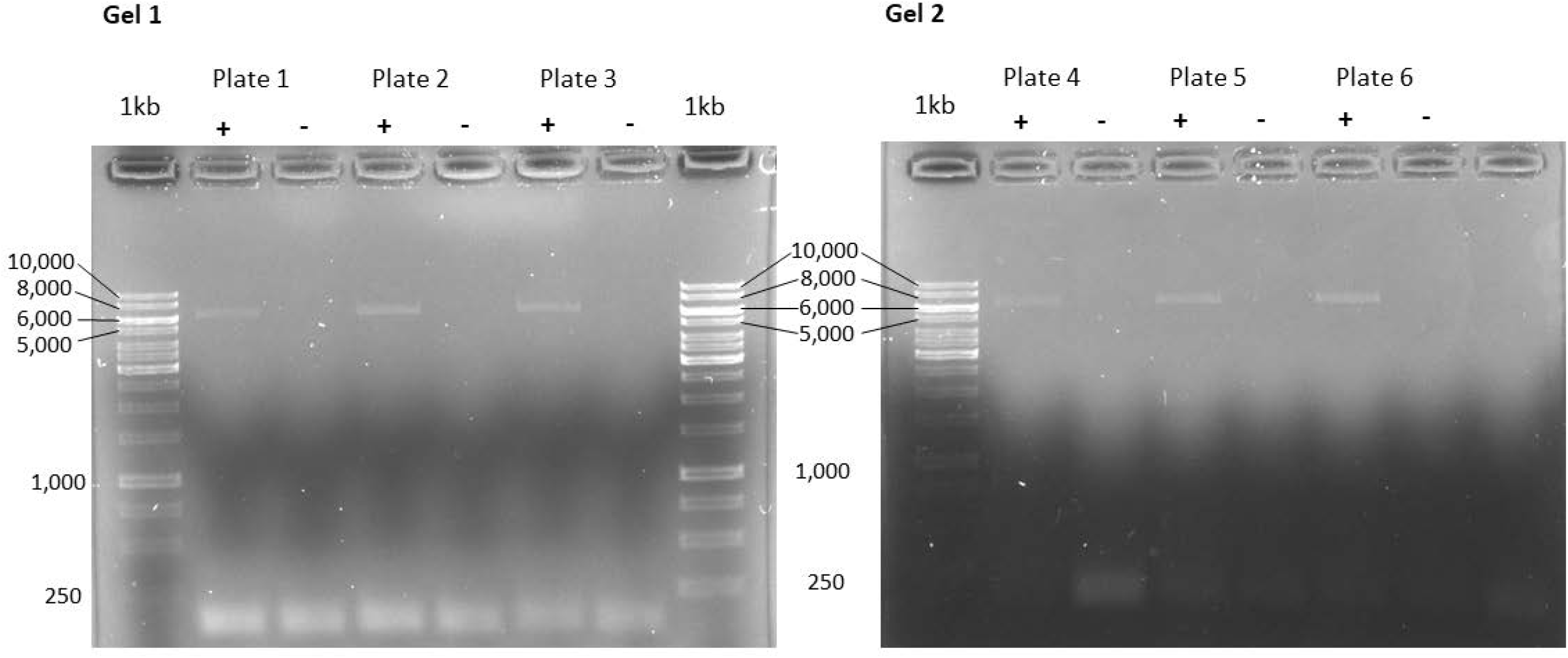
The plate setup (see Table S3) was repeated six times. Positive and negative control was included on each plate (6 plates in total). The controls were run on a 1 % TAE agarose gel shown here. We had successful amplification of correct size (6296 bp) in the positive controls (gel 1 and 2 (+): lane 2, 4, and 6) and no significant background contamination in the negative controls (gel 1 and 2 (-): lane 3, 5, and 7). GeneRuler 1kb ladder (ThermoFisher Scientific, Waltham, MA) is in lanes 1 and 8 (1kb) in gel 1 and lane 1 (1kb) in gel 2. (TIFF)

**Table S1.** This table provides the identified 81 nsSNPs (listed using the recommended format from the Human Genome Variation Society [83]) and details about their properties. The number of occurrences in the Vg variants are listed in column C. The scores from the substitution matrix (BLOSUM62) are listed for each nsSNP (Column D). Negative scores indicate that the physicochemical properties are not preserved. The calculated relative solvent accessible surface area (rASA) for each substituted residue position (column E) shows the percentage of the residue exposed to the solvent. Below 20% was considered buried. (XLSX)

**Table S2.** This table provides the identified 121 Vg variants. The Vg variants are numerate, and the table includes the identified nsSNPs per Vg variant. The nsSNPs are written using the same format as in Table S1. (XLSX)

**Table S3.** This table provides the PCR primers used for amplification of the *vg*-gene. Oligo sequence, melting temperature, and oligo size are provided for the forward and reverse primers. The forward and reverse primers were barcoded, creating 8 forward primers (F1 to F8) and 12 reverse primers (R1 to R12). Here we list the full oligo sequence, where the barcodes are written in red. The oligo size includes the primers, barcodes, and the Vg gene. We also include the plate setup, which shows the barcoded forward and reverse primer combinations used in each well. We repeated this setup for all 6 plates (see “Materials and Methods” for more details on the PCR protocol and Figure S1 for the complete workflow). (XLSX)

