## Supplementary material for "Identification of 121 variants of honey bee Vitellogenin protein sequences with structural differences at functional sites": Table S1

**Table S1. The identifed nsSNPs**

| Number | nsSNPs | Number of occurences on Vg variants | Modest (positive values) or drastic (negative values) | Exposed (>20 %), otherwise buried |
|---|---|---|---|---|
| 1 | p.Ala60Thr | 2 | 0 | 2 |
| 2 | p.Pro106Ser | 1 | -1 | 15 |
| 3 | p.Pro126Leu | 1 | -3 | 28 |
| 4 | p.Ile132Met | 16 | 1 | 1 |
| 5 | p.Gly146Ser | 3 | 0 | 16 |
| 6 | p.Thr305Met | 3 | -1 | 22 |
| 7 | p.Asn326Ser | 8 | 1 | 3 |
| 8 | p.His412Tyr | 3 | 2 | 3 |
| 9 | p.Val419Met | 3 | 1 | 5 |
| 10 | p.Ser467Asn | 1 | 1 | 34 |
| 11 | p.Ser471Asn | 3 | 1 | 59 |
| 12 | p.Ile489Val | 119 | 3 | 36 |
| 13 | p.Ala509Thr | 4 | 0 | 2 |
| 14 | p.Thr522Ile | 5 | -1 | 1 |
| 15 | p.Met525Ile | 1 | 1 | 61 |
| 16 | p.Asn560His | 1 | 1 | 77 |
| 17 | p.Thr594Met | 1 | -1 | 2 |
| 18 | p.Leu606Phe | 1 | 0 | 4 |
| 19 | p.Asp608Glu | 4 | 2 | 68 |
| 20 | p.Asp626Asn | 2 | 1 | 27 |
| 21 | p.Met639Ile | 4 | 1 | 6 |
| 22 | p.Ile640Val | 2 | 3 | 0 |
| 23 | p.Glu642Lys | 2 | 1 | 9 |
| 24 | p.Val661Met | 1 | 1 | 26 |
| 25 | p.Ser803Asn | 12 | 1 | 16 |
| 26 | p.Leu828Ile | 1 | 2 | 7 |
| 27 | p.Val834Met | 1 | 1 | 1 |
| 28 | p.Pro866Ser | 1 | -1 | 18 |
| 29 | p.Glu906Lys | 1 | 1 | 49 |
| 30 | p.Thr939Met | 1 | -1 | 18 |
| 31 | p.Val942Ala | 1 | 0 | 3 |
| 32 | p.Ile943Met | 1 | 1 | 0 |
| 33 | p.Pro958Ser | 1 | -1 | 91 |
| 34 | p.Arg969His | 1 | 0 | 27 |
| 35 | p.Thr984Met | 1 | -1 | 0 |
| 36 | p.Thr1013Met | 2 | -1 | 46 |
| 37 | p.Gly1016Asp | 1 | -1 | 29 |
| 38 | p.Gly1016Ser | 2 | 0 | 29 |
| 39 | p.Leu1072Phe | 1 | 0 | 31 |
| 40 | p.Asp1103Tyr | 1 | -3 | 65 |
| 41 | p.Thr1110Ala | 1 | 0 | 23 |
| 42 | p.Thr1110Ser | 14 | 1 | 23 |

| | | | | |
|---|---|---|---|---|
| 43 | p.Met1159Ile | 5 | 1 | 60 |
| 44 | p.Lys1171Asn | 4 | 0 | 61 |
| 45 | p.Arg1174Lys | 12 | 2 | 4 |
| 46 | p.Val1193Leu | 1 | 1 | 31 |
| 47 | p.Val1199Ile | 1 | 3 | 33 |
| 48 | p.Thr1207Ile | 1 | -1 | 23 |
| 49 | p.Ser1220Asn | 43 | 1 | 3 |
| 50 | p.Ser1235Arg | 2 | -1 | 58 |
| 51 | p.Ala1237Val | 1 | 0 | 35 |
| 52 | p.Arg1284Lys | 54 | 2 | 39 |
| 53 | p.Leu1291Ile | 22 | 2 | 17 |
| 54 | p.Arg1292Ser | 25 | -1 | 49 |
| 55 | p.Gly1302Glu | 2 | -2 | 35 |
| 56 | p.Val1311Met | 2 | 1 | 24 |
| 57 | p.Phe1357Val | 1 | -1 | 43 |
| 58 | p.Arg1385Lys | 1 | 2 | 37 |
| 59 | p.Ala1386Val | 1 | 0 | 16 |
| 60 | p.Val1397Met | 2 | 1 | 19 |
| 61 | p.Ile1398Val | 64 | 3 | 27 |
| 62 | p.Ala1451Val | 31 | 0 | 0 |
| 63 | p.Asp1491Asn | 3 | 1 | 43 |
| 64 | p.Ala1503Thr | 28 | 0 | 64 |
| 65 | p.Gly1504Arg | 1 | -2 | 11 |
| 66 | p.Val1508Met | 1 | 1 | 0 |
| 67 | p.Ile1536Val | 83 | 3 | 7 |
| 68 | p.Met1559Ile | 1 | 1 | 3 |
| 69 | p.Gly1565Ser | 1 | 0 | 0 |
| 70 | p.Thr1567Met | 1 | -1 | 3 |
| 71 | p.Thr1567Lys | 1 | -1 | 3 |
| 72 | p.Ser1587Phe | 1 | -1 | 32 |
| 73 | p.Ser1605Leu | 1 | -2 | 13 |
| 74 | p.Pro1620Ser | 1 | -1 | 19 |
| 75 | p.His1648Tyr | 1 | 2 | 42 |
| 76 | p.Leu1670Ser | 3 | -2 | 88 |
| 77 | p.Thr1676Ser | 31 | 1 | 76 |
| 78 | p.Gly1678Ser | 5 | 0 | 64 |
| 79 | p.Thr1692Ile | 1 | -1 | 2 |
| 80 | p.Ser1713Phe | 1 | -2 | 94 |
| 81 | p.Pro1755Ser | 1 | -1 | 72 |
