## Supplementary material for "Identification of 121 variants of honey bee Vitellogenin protein sequences with structural differences at functional sites": Table S2

**Table S2. The 121 Vg variants**

| Vg variants | nsSNP | | | | | | | | | | | |
|---|---|---|---|---|---|---|---|---|---|---|---|---|
| 1 | p.Ala60Thr | p.Ile489Val | p.Thr522Ile | p.Thr984Met | p.Lys1171Asn | p.Ser1220Asn | p.Arg1284Lys | p.Val1397Met | p.Ala1451Val | p.Ile1536Val | | |
| 2 | p.Ala60Thr | p.Ile489Val | p.Lys1171Asn | p.Ser1220Asn | p.Ser1235Arg | p.Val1397Met | p.Ile1536Val | | | | | |
| 3 | p.Pro106Ser | p.Ile489Val | p.Arg1284Lys | p.Ile1398Val | p.Ile1536Val | p.Thr1676Ser | | | | | | |
| 4 | p.Pro126Leu | p.Ile489Val | p.Ser1220Asn | p.Ala1451Val | | | | | | | | |
| 5 | p.Ile132Met | p.Asn326Ser | p.Ile489Val | p.Thr522Ile | p.Asp608Glu | p.Met639Ile | p.Lys1171Asn | p.Ser1220Asn | p.Arg1284Lys | p.Ile1398Val | p.Ala1451Val | p.Ile1536Val |
| 6 | p.Ile132Met | p.Ile489Val | p.Ser803Asn | p.Pro958Ser | p.Arg1174Lys | p.Leu1291Ile | p.Arg1292Ser | p.Val1508Met | p.Ile1536Val | p.Thr1676Ser | | |
| 7 | p.Ile132Met | p.Ile489Val | p.Ser803Asn | p.Arg1174Lys | p.Arg1284Lys | p.Ile1398Val | p.Ile1536Val | | | | | |
| 8 | p.Ile132Met | p.Ile489Val | p.Ser803Asn | p.Leu1291Ile | p.Arg1292Ser | p.Ile1398Val | p.Ile1536Val | | | | | |
| 9 | p.Ile132Met | p.Ile489Val | p.Ser803Asn | p.Leu1291Ile | p.Arg1292Ser | p.Ile1536Val | | | | | | |
| 10 | p.Ile132Met | p.Ile489Val | p.Val834Met | p.Arg1284Lys | p.Ile1398Val | p.Ile1536Val | p.Thr1676Ser | | | | | |
| 11 | p.Ile132Met | p.Ile489Val | p.Gly1016Asp | p.Ser1220Asn | p.Arg1284Lys | p.Ile1398Val | p.Ile1536Val | p.Thr1676Ser | p.Gly1678Ser | | | |
| 12 | p.Ile132Met | p.Ile489Val | p.Met1159Ile | p.Leu1291Ile | p.Arg1292Ser | p.Ile1398Val | p.Ile1536Val | | | | | |
| 13 | p.Ile132Met | p.Ile489Val | p.Arg1174Lys | p.Arg1284Lys | p.Ile1398Val | p.Ile1536Val | | | | | | |
| 14 | p.Ile132Met | p.Ile489Val | p.Arg1174Lys | p.Arg1284Lys | p.Ile1536Val | | | | | | | |
| 15 | p.Ile132Met | p.Ile489Val | p.Arg1174Lys | p.Leu1291Ile | p.Arg1292Ser | p.Ile1398Val | p.Ile1536Val | | | | | |
| 16 | p.Ile132Met | p.Ile489Val | p.Arg1284Lys | p.Ile1398Val | p.Ile1536Val | | | | | | | |
| 17 | p.Ile132Met | p.Ile489Val | p.Arg1284Lys | p.Ile1398Val | p.Ile1536Val | p.Thr1567Lys | p.Thr1676Ser | p.Gly1678Ser | | | | |
| 18 | p.Ile132Met | p.Ile489Val | p.Arg1284Lys | p.Ile1536Val | p.Thr1676Ser | | | | | | | |
| 19 | p.Ile132Met | p.Ile489Val | p.Ile1398Val | p.Asp1491Asn | p.Ile1536Val | | | | | | | |
| 20 | p.Ile132Met | p.Ile489Val | p.Ile1536Val | | | | | | | | | |
| 21 | p.Gly146Ser | p.Ile489Val | p.Thr1110Ser | p.Ser1220Asn | p.Ala1451Val | p.Ala1503Thr | | | | | | |
| 22 | p.Gly146Ser | p.Ile489Val | p.Val1199Ile | p.Arg1284Lys | p.Ile1398Val | p.Ile1536Val | | | | | | |
| 23 | p.Gly146Ser | p.Ile489Val | p.Arg1284Lys | p.Ala1451Val | p.Ala1503Thr | | | | | | | |
| 24 | p.Thr305Met | p.His412Tyr | p.Val419Met | p.Ser471Asn | p.Ile489Val | p.Thr522Ile | p.Asp608Glu | p.Met639Ile | p.Lys1171Asn | p.Ser1220Asn | p.Ala1451Val | p.Ala1503Thr |
| 25 | p.Thr305Met | p.His412Tyr | p.Val419Met | p.Ser471Asn | p.Ile489Val | p.Thr522Ile | p.Asp608Glu | p.Met639Ile | p.Arg1284Lys | p.Ile1398Val | p.Ile1536Val | |
| 26 | p.Thr305Met | p.Ile489Val | p.Thr1110Ser | p.Ser1220Asn | p.Leu1291Ile | p.Arg1292Ser | p.Ala1451Val | p.Ala1503Thr | | | | |
| 27 | p.Asn326Ser | p.Ile489Val | p.Ile640Val | p.Gly1504Arg | p.Leu1670Ser | | | | | | | |
| 28 | p.Asn326Ser | p.Ile489Val | p.Ile640Val | p.Leu1670Ser | | | | | | | | |
| 29 | p.Asn326Ser | p.Ile489Val | p.Thr1110Ser | p.Ser1220Asn | p.Leu1291Ile | p.Arg1292Ser | p.Ala1503Thr | | | | | |
| 30 | p.Asn326Ser | p.Ile489Val | p.Ser1220Asn | | | | | | | | | |
| 31 | p.Asn326Ser | p.Ile489Val | p.Ser1220Asn | p.Ala1451Val | | | | | | | | |
| 32 | p.Asn326Ser | p.Ile489Val | p.Ser1220Asn | p.Ala1451Val | p.Ala1503Thr | | | | | | | |
| 33 | p.Asn326Ser | p.Ile489Val | p.Ser1220Asn | p.Ala1451Val | p.Ser1605Leu | | | | | | | |
| 34 | p.His412Tyr | p.Val419Met | p.Ser467Asn | p.Ile489Val | p.Asn560His | p.Thr1110Ala | p.Ser1235Arg | p.Val1311Met | p.Ile1536Val | | | |
| 35 | p.Ser471Asn | p.Ile489Val | p.Thr522Ile | p.Asp608Glu | p.Met639Ile | p.Arg1284Lys | p.Ile1398Val | p.Ile1536Val | | | | |
| 36 | p.Ile489Val | p.Ala509Thr | p.Arg1284Lys | p.Gly1302Glu | p.Ile1398Val | p.Ile1536Val | | | | | | |
| 37 | p.Ile489Val | p.Ala509Thr | p.Arg1284Lys | p.Ile1398Val | p.Ile1536Val | | | | | | | |
| 38 | p.Ile489Val | p.Ala509Thr | p.Arg1284Lys | p.Ile1398Val | p.Ile1536Val | p.Thr1676Ser | | | | | | |
| 39 | p.Ile489Val | p.Ala509Thr | p.Arg1284Lys | p.Ile1536Val | p.Thr1676Ser | p.Ser1713Phe | | | | | | |
| 40 | p.Ile489Val | p.Met525Ile | p.Ser1220Asn | p.Ala1451Val | p.Ala1503Thr | p.Ile1536Val | | | | | | |
| 41 | p.Ile489Val | p.Thr594Met | p.Ser1220Asn | | | | | | | | | |
| 42 | p.Ile489Val | p.Leu606Phe | p.Arg1284Lys | p.Ile1398Val | p.Ile1536Val | p.Thr1676Ser | | | | | | |
| 43 | p.Ile489Val | p.Asp626Asn | p.Arg1284Lys | p.Ile1398Val | p.Ile1536Val | | | | | | | |
| 44 | p.Ile489Val | p.Asp626Asn | p.Arg1284Lys | p.Ile1398Val | p.Ile1536Val | p.Thr1676Ser | | | | | | |
| 45 | p.Ile489Val | p.Glu642Lys | p.Arg1284Lys | p.Ile1398Val | p.Ile1536Val | p.Thr1676Ser | | | | | | |
| 46 | p.Ile489Val | p.Glu642Lys | p.Arg1284Lys | p.Ala1451Val | p.Ala1503Thr | | | | | | | |
| 47 | p.Ile489Val | p.Val661Met | p.Ser1220Asn | p.Ala1451Val | p.Ala1503Thr | | | | | | | |
| 48 | p.Ile489Val | p.Ser803Asn | p.Met1159Ile | p.Arg1284Lys | p.Ile1398Val | p.Asp1491Asn | p.Ile1536Val | p.Thr1676Ser | | | | |
| 49 | p.Ile489Val | p.Ser803Asn | p.Met1159Ile | p.Leu1291Ile | p.Arg1292Ser | p.Ile1398Val | p.Ile1536Val | | | | | |
| 50 | p.Ile489Val | p.Ser803Asn | p.Arg1284Lys | p.Ile1398Val | p.Ala1503Thr | p.Ile1536Val | | | | | | |
| 51 | p.Ile489Val | p.Ser803Asn | p.Arg1284Lys | p.Ile1398Val | p.Ile1536Val | p.Thr1676Ser | | | | | | |
| 52 | p.Ile489Val | p.Ser803Asn | p.Leu1291Ile | p.Arg1292Ser | p.Ile1536Val | | | | | | | |
| 53 | p.Ile489Val | p.Ser803Asn | p.Ile1398Val | p.Asp1491Asn | p.Ile1536Val | | | | | | | |
| 54 | p.Ile489Val | p.Ser803Asn | p.Ile1398Val | p.Ile1536Val | p.Thr1676Ser | p.Gly1678Ser | | | | | | |
| 55 | p.Ile489Val | p.Ser803Asn | p.Ser1220Asn | p.Arg1284Lys | p.Ile1398Val | p.Ile1536Val | p.Pro1755Ser | | | | | |
| 56 | p.Ile489Val | p.Leu828Ile | p.Arg1284Lys | p.Ile1398Val | p.Ile1536Val | p.Thr1676Ser | p.Gly1678Ser | | | | | |
| 57 | p.Ile489Val | p.Pro866Ser | p.Thr1110Ser | p.Ser1220Asn | p.Ala1451Val | p.Ala1503Thr | | | | | | |
| 58 | p.Ile489Val | p.Glu906Lys | p.Ser1220Asn | p.Ala1451Val | p.Ala1503Thr | | | | | | | |
| 59 | p.Ile489Val | p.Thr939Met | p.Arg1284Lys | p.Ile1398Val | p.Ile1536Val | p.Thr1676Ser | | | | | | |
| 60 | p.Ile489Val | p.Val942Ala | p.Ser1220Asn | | | | | | | | | |
| 61 | p.Ile489Val | p.Ile943Met | p.Arg1284Lys | p.Ile1398Val | p.Ile1536Val | | | | | | | |
| 62 | p.Ile489Val | p.Arg969His | p.Arg1284Lys | p.Ile1398Val | p.Ile1536Val | p.Thr1676Ser | | | | | | |
| 63 | p.Ile489Val | p.Thr1013Met | p.Arg1284Lys | p.Ile1398Val | p.Ile1536Val | | | | | | | |
| 64 | p.Ile489Val | p.Thr1013Met | p.Arg1284Lys | p.Ile1398Val | p.Ile1536Val | p.Thr1676Ser | | | | | | |
| 65 | p.Ile489Val | p.Gly1016Ser | p.Arg1174Lys | p.Leu1291Ile | p.Arg1292Ser | p.Ile1398Val | p.Ile1536Val | | | | | |
| 66 | p.Ile489Val | p.Gly1016Ser | p.Arg1284Lys | p.Ile1398Val | p.Ile1536Val | | | | | | | |
| 67 | p.Ile489Val | p.Leu1072Phe | p.Thr1110Ser | p.Ser1220Asn | p.Ala1451Val | p.Ala1503Thr | | | | | | |
| 68 | p.Ile489Val | p.Asp1103Tyr | p.Thr1110Ser | p.Ser1220Asn | p.Ala1451Val | p.Ala1503Thr | | | | | | |
| 69 | p.Ile489Val | p.Thr1110Ser | p.Ser1220Asn | | | | | | | | | |
| 70 | p.Ile489Val | p.Thr1110Ser | p.Ser1220Asn | p.Leu1291Ile | p.Arg1292Ser | p.Ala1451Val | p.Ala1503Thr | | | | | |
| 71 | p.Ile489Val | p.Thr1110Ser | p.Ser1220Asn | p.Ile1398Val | p.Ile1536Val | | | | | | | |
| 72 | p.Ile489Val | p.Thr1110Ser | p.Ser1220Asn | p.Ala1451Val | | | | | | | | |
| 73 | p.Ile489Val | p.Thr1110Ser | p.Ser1220Asn | p.Ala1451Val | p.Ala1503Thr | | | | | | | |
| 74 | p.Ile489Val | p.Thr1110Ser | p.Ser1220Asn | p.Ala1503Thr | | | | | | | | |
| 75 | p.Ile489Val | p.Thr1110Ser | p.Ser1220Asn | p.Ile1536Val | p.Thr1692Ile | | | | | | | |
| 76 | p.Ile489Val | p.Thr1110Ser | p.Arg1284Lys | p.Ile1398Val | p.Ala1503Thr | p.Ile1536Val | | | | | | |
| 77 | p.Ile489Val | p.Met1159Ile | p.Leu1291Ile | p.Arg1292Ser | p.Ile1398Val | p.Ile1536Val | | | | | | |
| 78 | p.Ile489Val | p.Met1159Ile | p.Ile1398Val | p.Ile1536Val | | | | | | | | |
| 79 | p.Ile489Val | p.Arg1174Lys | p.Arg1284Lys | p.Ile1398Val | p.Ile1536Val | | | | | | | |
| 80 | p.Ile489Val | p.Arg1174Lys | p.Arg1284Lys | p.Ile1398Val | p.Ile1536Val | p.Thr1676Ser | | | | | | |
| 81 | p.Ile489Val | p.Arg1174Lys | p.Arg1284Lys | p.Ile1536Val | | | | | | | | |

| | | | | | | | | |
|---|---|---|---|---|---|---|---|---|
| 82 | p.Ile489Val | p.Arg1174Lys | p.Arg1284Lys | p.Ile1536Val | p.Thr1676Ser | | | |
| 83 | p.Ile489Val | p.Arg1174Lys | p.Leu1291Ile | p.Arg1292Ser | p.Ile1398Val | p.Ile1536Val | | |
| 84 | p.Ile489Val | p.Arg1174Lys | p.Leu1291Ile | p.Arg1292Ser | p.Ile1398Val | p.Ile1536Val | p.Met1559Ile | |
| 85 | p.Ile489Val | p.Val1193Leu | p.Arg1385Lys | p.Ile1398Val | p.Pro1620Ser | | | |
| 86 | p.Ile489Val | p.Thr1207Ile | p.Arg1284Lys | p.Ile1398Val | p.Ile1536Val | p.Thr1676Ser | | |
| 87 | p.Ile489Val | p.Ser1220Asn | p.Ala1237Val | p.Ala1451Val | p.Ala1503Thr | | | |
| 88 | p.Ile489Val | p.Ser1220Asn | p.Arg1284Lys | p.Ile1398Val | p.Ile1536Val | | | |
| 89 | p.Ile489Val | p.Ser1220Asn | p.Leu1291Ile | p.Arg1292Ser | | | | |
| 90 | p.Ile489Val | p.Ser1220Asn | p.Leu1291Ile | p.Arg1292Ser | p.Ala1451Val | | | |
| 91 | p.Ile489Val | p.Ser1220Asn | p.Leu1291Ile | p.Arg1292Ser | p.Ala1451Val | p.Ala1503Thr | | |
| 92 | p.Ile489Val | p.Ser1220Asn | p.Leu1291Ile | p.Arg1292Ser | p.Ala1451Val | p.Ala1503Thr | p.Thr1567Met | |
| 93 | p.Ile489Val | p.Ser1220Asn | p.Arg1292Ser | | | | | |
| 94 | p.Ile489Val | p.Ser1220Asn | p.Arg1292Ser | p.Ile1536Val | | | | |
| 95 | p.Ile489Val | p.Ser1220Asn | p.Ile1398Val | p.Ile1536Val | p.Gly1565Ser | p.Thr1676Ser | | |
| 96 | p.Ile489Val | p.Ser1220Asn | p.Ala1451Val | | | | | |
| 97 | p.Ile489Val | p.Ser1220Asn | p.Ala1451Val | p.Ala1503Thr | | | | |
| 98 | p.Ile489Val | p.Ser1220Asn | p.Ala1451Val | p.Ala1503Thr | p.Ile1536Val | | | |
| 99 | p.Ile489Val | p.Ser1220Asn | p.Ala1451Val | p.Ala1503Thr | p.Thr1676Ser | | | |
| 100 | p.Ile489Val | p.Ser1220Asn | p.Ala1451Val | p.Ile1536Val | | | | |
| 101 | p.Ile489Val | p.Arg1284Lys | p.Gly1302Glu | p.Phe1357Val | p.Ile1398Val | p.Ile1536Val | | |
| 102 | p.Ile489Val | p.Arg1284Lys | p.Val1311Met | p.Ile1398Val | p.Ile1536Val | p.Thr1676Ser | | |
| 103 | p.Ile489Val | p.Arg1284Lys | p.Ala1386Val | p.Ile1398Val | p.Ile1536Val | p.Ser1587Phe | p.His1648Tyr | p.Thr1676Ser |
| 104 | p.Ile489Val | p.Arg1284Lys | p.Ile1398Val | | | | | |
| 105 | p.Ile489Val | p.Arg1284Lys | p.Ile1398Val | p.Ala1503Thr | p.Ile1536Val | | | |
| 106 | p.Ile489Val | p.Arg1284Lys | p.Ile1398Val | p.Ala1503Thr | p.Ile1536Val | p.Thr1676Ser | | |
| 107 | p.Ile489Val | p.Arg1284Lys | p.Ile1398Val | p.Ile1536Val | | | | |
| 108 | p.Ile489Val | p.Arg1284Lys | p.Ile1398Val | p.Ile1536Val | p.Thr1676Ser | | | |
| 109 | p.Ile489Val | p.Arg1284Lys | p.Ile1398Val | p.Ile1536Val | p.Thr1676Ser | p.Gly1678Ser | | |
| 110 | p.Ile489Val | p.Arg1284Lys | p.Ala1451Val | p.Ala1503Thr | | | | |
| 111 | p.Ile489Val | p.Arg1284Lys | p.Ile1536Val | | | | | |
| 112 | p.Ile489Val | p.Arg1284Lys | p.Ile1536Val | p.Thr1676Ser | | | | |
| 113 | p.Ile489Val | p.Leu1291Ile | p.Arg1292Ser | p.Ile1398Val | p.Ile1536Val | | | |
| 114 | p.Ile489Val | p.Leu1291Ile | p.Arg1292Ser | p.Ile1398Val | p.Ile1536Val | p.Thr1676Ser | | |
| 115 | p.Ile489Val | p.Leu1291Ile | p.Arg1292Ser | p.Ile1536Val | | | | |
| 116 | p.Ile489Val | p.Leu1291Ile | p.Arg1292Ser | p.Ile1536Val | p.Thr1676Ser | | | |
| 117 | p.Ile489Val | p.Arg1292Ser | p.Ile1398Val | p.Ile1536Val | | | | |
| 118 | p.Ile489Val | p.Ile1398Val | p.Ile1536Val | | | | | |
| 119 | p.Ile489Val | p.Ala1451Val | p.Ala1503Thr | | | | | |
| 120 | p.Ile1398Val | p.Ile1536Val | | | | | | |
| 121 | p.Ile1398Val | p.Ile1536Val | p.Leu1670Ser | | | | | |
