## Supplementary material for "Identification of 121 variants of honey bee Vitellogenin protein sequences with structural differences at functional sites": Table S3

**Table S3. PCR primers, barcodes and PCR plate setup**

| Primers for *vg* gene | Oligo sequence (5' to 3') | Tm | Oligo size |
|---|---|---|---|
| Forward | AGCCGAATCAAATGCATCGT | 58.6 | 20 |
| Reverse | ACGAAAGAAAGGATTATTGAAAACA | 56 | 25 |

| Primer and Barcodes | Oligo sequence (5' to 3') | Oligo size | Full-length fragment size |
|---|---|---|---|
| F1 | AAGAAAGTTGTCGGTGTCTTTGTGAGCCGAATCAAATGCATCGT | 44 | 6296 bp |
| F2 | TCGATTCCGTTTGTAGTCGTCTGTAGCCGAATCAAATGCATCGT | 44 | 6296 bp |
| F3 | GAGTCTTGTGTCCCAGTTACCAGGAGCCGAATCAAATGCATCGT | 44 | 6296 bp |
| F4 | TTCGGATTCTATCGTGTTTCCCTAAGCCGAATCAAATGCATCGT | 44 | 6296 bp |
| F5 | CTTGTCCAGGGTTTGTGTAACCTTAGCCGAATCAAATGCATCGT | 44 | 6296 bp |
| F6 | TTCTCGCAAAGGCAGAAAGTAGTCAGCCGAATCAAATGCATCGT | 44 | 6296 bp |
| F7 | GTGTTACCGTGGGAATGAATCCTTAGCCGAATCAAATGCATCGT | 44 | 6296 bp |
| F8 | TTCAGGGAACAAACCAAGTTACGTAGCCGAATCAAATGCATCGT | 44 | 6296 bp |
| R1 | AGAACGACTTCCATACTCGTGTGAACGAAAGAAAGGATTATTGAAAACA | 49 | 6296 bp |
| R2 | AACGAGTCTCTTGGGACCCATAGAACGAAAGAAAGGATTATTGAAAACA | 49 | 6296 bp |
| R3 | AGGTCTACCTCGCTAACACCACTGACGAAAGAAAGGATTATTGAAAACA | 49 | 6296 bp |
| R4 | CGTCAACTGACAGTGGTTCGTACTACGAAAGAAAGGATTATTGAAAACA | 49 | 6296 bp |
| R5 | ACCCTCCAGGAAAGTACCTCTGATACGAAAGAAAGGATTATTGAAAACA | 49 | 6296 bp |
| R6 | CCAAACCCAACAACCTAGATAGGCACGAAAGAAAGGATTATTGAAAACA | 49 | 6296 bp |
| R7 | GTTCCTCGTGCAGTGTCAAGAGATACGAAAGAAAGGATTATTGAAAACA | 49 | 6296 bp |
| R8 | TTGCGTCCTGTTACGAGAACTCATACGAAAGAAAGGATTATTGAAAACA | 49 | 6296 bp |
| R9 | GAGCCTCTCATTGTCCGTTCTCTAACGAAAGAAAGGATTATTGAAAACA | 49 | 6296 bp |
| R10 | ACCACTGCCATGTATCAAAGTACGACGAAAGAAAGGATTATTGAAAACA | 49 | 6296 bp |
| R11 | CTTACTACCCAGTGAACCTCCTCGACGAAAGAAAGGATTATTGAAAACA | 49 | 6296 bp |
| R12 | GCATAGTTCTGCATGATGGGTTAGACGAAAGAAAGGATTATTGAAAACA | 49 | 6296 bp |

**Plate setup**

| | 1 | 2 | 3 | 4 | 5 | 6 | 7 | 8 | 9 | 10 | 11 | 12 |
|---|---|---|---|---|---|---|---|---|---|---|---|---|
| A | F1R1 | F1R2 | F1R3 | F1R4 | F1R5 | F1R6 | F1R7 | F1R8 | F1R9 | F1R10 | F1R11 | F1R12 |
| B | F2R1 | F2R2 | F2R3 | F2R4 | F2R5 | F2R6 | F2R7 | F2R8 | F2R9 | F2R10 | F2R11 | F2R12 |
| C | F3R1 | F3R2 | F3R3 | F3R4 | F3R5 | F3R6 | F3R7 | F3R8 | F3R9 | F3R10 | F3R11 | F3R12 |
| D | F4R1 | F4R2 | F4R3 | F4R4 | F4R5 | F4R6 | F4R7 | F4R8 | F4R9 | F4R10 | F4R11 | F4R12 |
| E | F5R1 | F5R2 | F5R3 | F5R4 | F5R5 | F5R6 | F5R7 | F5R8 | F5R9 | F5R10 | F5R11 | F5R12 |
| F | F6R1 | F6R2 | F6R3 | F6R4 | F6R5 | F6R6 | F6R7 | F6R8 | F6R9 | F6R10 | F6R11 | F6R12 |
| G | F7R1 | F7R2 | F7R3 | F7R4 | F7R5 | F7R6 | F7R7 | F7R8 | F7R9 | F7R10 | F7R11 | F7R12 |
| H | F8R1 | F8R2 | F8R3 | F8R4 | F8R5 | F8R6 | F8R7 | F8R8 | F8R9 | F8R10 | F8R11 | F8R12 |
